# FMO4 drives lung adenocarcinoma by stabilizing the MAT2A/MAT2B complex and hindering ferroptosis

**DOI:** 10.1101/2025.03.31.646284

**Authors:** David Bracquemond, Sayyed Jalil Mahdizadeh, Lisa Brunet, Quentin-Dominique Thomas, Céline Bouclier, Marie Colomb, Sylvia Bourdel, Eric Fabbrizio, Hélène Tourriere, Laura Papon, Xavier Quantin, Alba Santos, Irene Ferrer, Sylvain Henry, Marie Grosgeorges, Valerio Perna, Maria D. Lozano, José Luis Solórzano, Jozef P. Bossowski, Thales Papagiannakopoulos, Marcello Tiseo, Roberta Minari, Andrei Turtoi, Alain Mangé, Luis M. Montuenga, Luis Paz-Ares, Julien Faget, Leif A. Eriksson, Maicol Mancini, Antonio Maraver

## Abstract

Lung cancer is the leading cause of death by cancer in the world and finding new targets is a major medical need to tackle this disease. Here, upon proteomic analysis to identify common players in oncogenic EGFR- and KRAS-driven lung adenocarcinoma mouse models, we uncovered a largely unknown protein in cancer, flavin-containing monooxygenase 4 (FMO4), whose expression was increased in lung tumors compared with adjacent lung tissue. FMO4 expression was strongly increased also in lung cancer samples from patients compared with healthy lung, and its expression level was inversely correlated with overall survival. Remarkably, in vivo deletion of FMO4 greatly decreased tumor burden and increased survival in oncogenic KRAS-driven lung adenocarcinoma mice unveiling its crucial role in tumor biology. Mechanistically, we found that FMO4 loss of function promotes ferroptosis and cooperates with ferroptosis inducers *in vitro* and *in vivo*. Moreover, FMO4 facilitates the interaction between MAT2A and MAT2B, promoting the generation of cysteine from methionine, which in turn boosts the generation of glutathione, thus protecting lung adenocarcinoma against ferroptosis. In summary we identified a new target in lung adenocarcinoma with important implications in cancer biology.

## INTRODUCTION

Despite intense research efforts translated into major clinical advances, lung cancer remains the most common cause of cancer-related death worldwide ^1^. Therefore, it is crucial to find new lung cancer vulnerabilities to broad the therapeutic options in this deadly disease.

Here, we uncovered the role in lung adenocarcinoma (accounting for around 50% of all lung cancer patients), of flavin-containing monooxygenase 4 (FMO4). This protein belongs to the FMO family that appear very early in the animal kingdom ^2^ and it is very well conserved throughout evolution ^3^. FMOs were discovered as detoxifiers of different xenobiotics ^4^. However, in recent years, several studies showed that FMOs are also implicated in genetic diseases (for instance, FMO3 in trimethylaminuria), in different metabolic disorders as diabetes, as well as in aging ^5^. In this sense, the expression of several FMOs is increased upon calorie restriction in several animal models ^5^. In accordance with this, overexpression of the *Caenorhabditis elegans* orthologue of human FMO2 is sufficient to increase the lifespan in these worms ^6^, indicating that these proteins play major roles in aging and protection to different types of stresses. Conversely, FMO putative roles in cancer, including lung are not well known. To the best of our knowledge, the only studies in lung cancer were focused mainly on FMO activity as carcinogen detoxifiers ^7^ or therapeutic drug modifiers ^8^. Therefore, their direct roles in lung cancer initiation and/or maintenance remained to be explored.

In this study we found that FMO4 was strongly expressed in clinical samples from patients with different lung cancer types including lung adenocarcinoma, the most prevalent subtype. Moreover, FMO4 expression was inversely correlated with survival in lung adenocarcinoma patients. Mechanistically, we showed that FMO4 expression was increased by acute oxidative stress and we identified aryl hydrocarbon receptor (AHR), a transcription factor that among others control the expression of the oxidative stress master regulator NRF2 ^9^, as the mediator of FMO4 expression. Interestingly, FMO4 loss of function sensitized cells against ferroptosis, a type of cellular death promoted by lipid peroxidation with increased interest in the cancer field, including lung cancer, as an attractive vulnerability to target ^10^. Still, we need to further increase our knowledge in this type of death to improve the current therapeutic options ^11^. Following this idea, we also found that FMO4 binds to and regulates methionine adenosyltransferase 2A (MAT2A), a rate-limiting enzyme of the methionine cycle ^12^ and likewise, FMO4 loss of function deregulate this metabolic pathway that catalyzes the conversion of methionine into s-adenosylmethionine (SAM), which in turn is converted into s-adenosylhomocysteine (SAH), and finally into homocysteine that can be regenerated into methionine to end the cycle, or enter into the transsulfuration pathway. This downstream connected pathway generates cysteine and reduced glutathione ^12^, which is crucial in the activity of glutathione peroxidase 4 (GPX4) to prevent ferroptosis ^13^. Finally, using AlphaFold, an artificial intelligence-based tool to predict protein structure and molecular dynamics simulations ^14^, we provided evidence that FMO4 facilitates MAT2A/MAT2B complex formation.

In summary, we uncovered that FMO4 is a novel key player in lung cancer with critical functions in controlling ferroptosis during lung adenocarcinoma development and homeostasis.

## RESULTS

### FMO4 is highly expressed in mouse and in human lung cancer

Here, we used two genetically engineered mouse models (GEMMs) of lung adenocarcinoma induced by lung-specific expression of oncogenic KRAS ^15^ with p53 conditional deletion (i.e., KRAS^G12V^/p53^flox/flox^ strain) and of oncogenic EGFR ^16^ (i.e., EGFR^T790M/L858R^/CCSPrtTA strain) to identify pathways that were affected in both strains. The proteomic analysis of lung cancer samples and adjacent healthy tissue from these two GEMMs identified 250 proteins that were upregulated or downregulated in lung cancer compared with normal lung tissue in both strains (Extended Data Table 1). Using the “STRING” tool, we found that “FMO oxidases nucleophiles” showed the highest strength in enrichment among the obtained Reactome pathways (Fig. 1a). These protein family is known to play important roles in different diseases ^5^; but conversely, its possible role in cancer is largely unknown. Hence, we focused our attention into this protein family to shed light into such putative function that could unveil new targets for lung cancer.

**Fig. 1.**
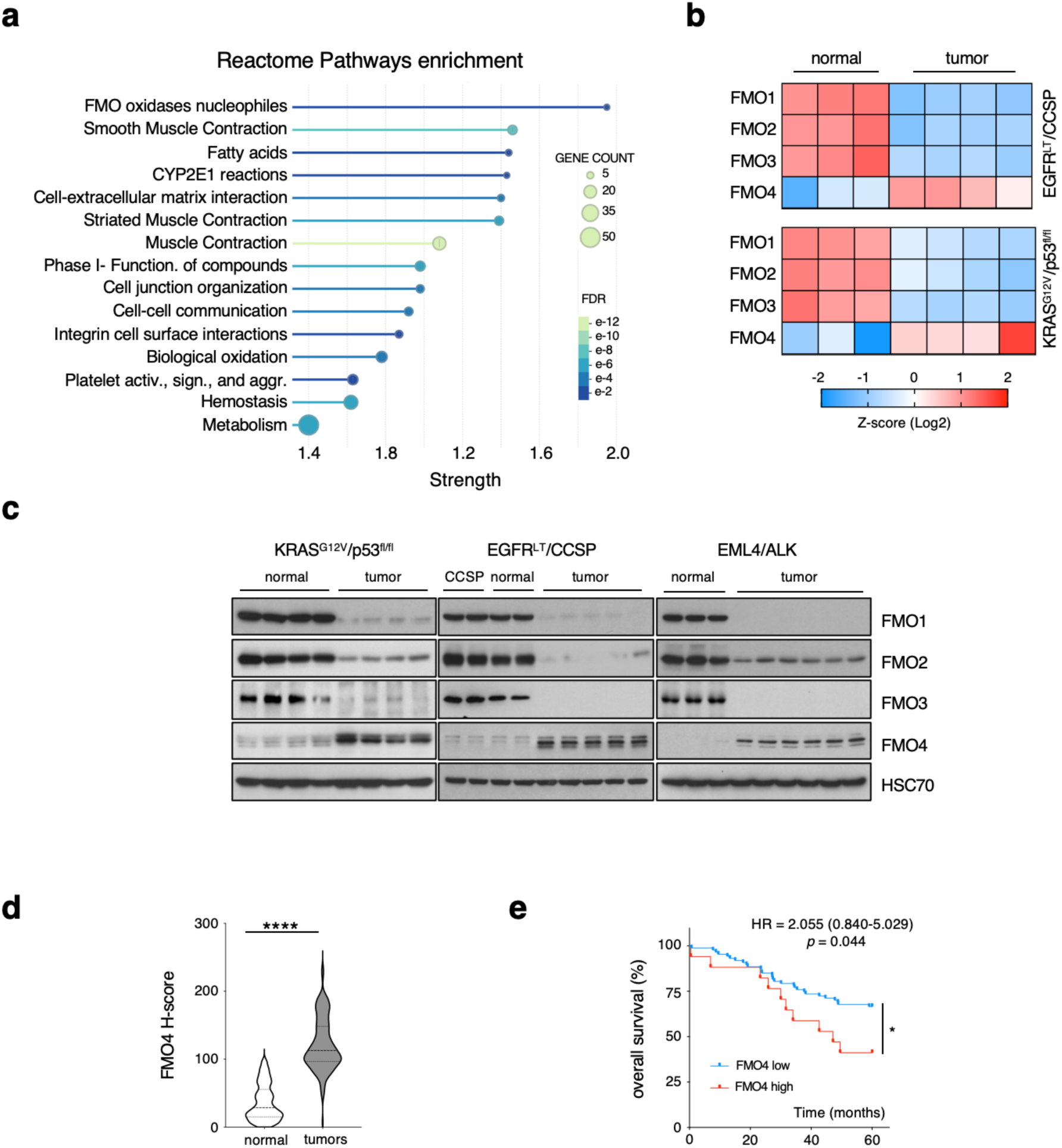
FMO4 is highly expressed in mouse and human lung cancer. **a)** REACTOME pathway enrichment analysis of commonly upregulated or downregulated proteins in lung cancer compared with healthy adjacent lung tissue from KRAS^G12V^/p53^flox/flox^ and EGFR^T790M/L858R^/CCSPrtTA mice. **b)** Heatmap of FMO1, 2, 3, 4 protein expression in lung tumors (n = 4/each) and healthy lung samples (n = 3/each) from EGFR^T790M/L858R^/CCSPrtTA and KRAS^G12V^/p53^flox/flox^ mice. **c**) Immunoblotting of the indicated proteins coming from (left panel) KRAS^G12V^/p53^floxflox^ mice (normal lung tissue, n = 4; tumors, n = 4), (middle panel) CCSP control mice (normal lung, n = 2) and EGFR^T790M/L858R^/CCSP mice (normal lung, n = 2; and tumors, n = 4), and (right panel) EML4/ALK mice (normal lung, n = 3; and tumors, n = 6). **d)** Violin plots showing the FMO4 H-score in normal lung (n = 244) and tumor (n = 185) biopsies. The median and upper/lower quartiles are depicted with dashed and dotted lines respectively; \*\*\*\**p* <0.0001, two-tailed unpaired Student’s *t*-test. **e**) Kaplan Meier curves showing the overall survival of patients with lung adenocarcinoma according to FMO4 expression (FMO4-low n = 88 and FMO4-high n = 18). Statistical significance was calculated with the Mantel-Cox log-rank test: \**p* = 0.044 [HR = 2.055 (95%CI = 0.840–5.029)].

In mice and humans, *FMO1* to *4* are clustered on chromosome 1. Conversely, mouse *Fmo5* is on chromosome 3 and human *FMO5* is on chromosome 1, but far away from the cluster ^5^, that is in accordance with the branching evolution of FMOs where FMO5 was the first diverging from the ancestor FMO and the only one having Baeyer– Villiger oxidation capacity in modern FMOs ^17^. Therefore, we performed an unsupervised clustering of FMO1 to 4 in the same samples subjected to proteomics above. Intriguingly, unlike FMO1, 2 and 3, FMO4 expression was decreased in healthy lung and increased in tumor samples in both cancer mouse models (Fig. 1b). To validate this interesting observation, we analyzed by western blotting another set of lung cancer samples from both GEMMs, and to extend our findings we included also another GEMM of lung adenocarcinoma induced by an endogenous EML4/ALK translocation ^18^. Interestingly, we observed the opposite expression patterns between FMO4 and FMO1, 2 and 3 in all GEMMs, confirming and expanding our original observation (Fig. 1c). Also, RNA expression analysis showed that *Fmo4* was upregulated in tumors from the three GEMMs compared with healthy lung, whereas *Fmo1*, *Fmo2*, *Fmo3* were downregulated (Extended Data Fig. 1a). Therefore, we conclude that FMO4 is a new tumor marker in murine lung adenocarcinoma induced by the three most prevalent oncogenic drivers in patients with human lung adenocarcinoma (∼50%) ^19^.

To increase the clinical relevance of our findings, we analyzed the FMO4 expression using the histoscore (H-score) ^20^ in a previously published tissue microarray (TMA) ^21^ that includes healthy lung tissue and non-small cell lung cancer (NSCLC) samples (adenocarcinoma, squamous cell carcinoma and other minor subtypes, Extended Data Table 2). FMO4 expression level was increased by more than 4 x-fold in tumor samples compared with controls (Fig. 1d, Extended Data Fig. 1b) and this was also true in all different tumor histological subtypes separately (Extended Data Fig. 1c).

Then, we analyzed *FMO4* copy number alterations in ∼2000 lung adenocarcinoma samples from five publicly available datasets in cBioPortal ^22–26^. In these datasets, *FMO4* genomic region was amplified in 3-8% of samples (Extended Data Fig. 1d). Conversely, we found deep deletions, which are associated with anti-tumoral functions, at this locus only in two datasets and at very low frequency (Extended Data Fig. 1d). As *FMO1* to *4* are in the same cluster on chromosome 1, it is not surprising that the copy number alteration frequency was similar in *FMO1*, *2*, *3* and *4* loci. However, comparison of *FMO1*, *FMO2*, *FMO3* and *FMO4* mRNA levels in patients with two copies (diploid) and with ≥3 copies (amplified) in each respective locus showed that only *FMO4* expression was increased in patients with amplification (Extended Data Fig. 1e). This indicates that the increased copy number affected only *FMO4* expression.

Next, we analyzed in the same lung cancer TMA ^21^, the correlation between FMO4 levels and 5-year overall survival. We found a trend (*p* = 0.097) towards shorter survival in patients with high FMO4 expression compared with low FMO4 expression (Extended Data Fig. 1f). Same analysis using a different TMA from another cohort of patients with only lung adenocarcinoma ^27^ (Extended Data Table 3) showed a very similar pattern but in this second cohort it was significant (HR 2.055, 95% CI 0.84-5.029 *p* = 0.044) (Fig. 1e).

These clinical data indicate that FMO4 could have a protumoral role in lung cancer and identified high FMO4 expression as a poor prognostic factor in lung adenocarcinoma patients.

### FMO4 is required for oncogenic KRAS-driven LUAD lung adenocarcinoma

To determine FMO4 role in lung tumorigenesis, we used a lentiviral shRNA technology ^28^ to constitutively express in the KRAS^G12D^/p53^flox/flox^ (KP) GEMM a non-targeting shRNA (*shNT*) or a shRNA against murine FMO4 (*shFmo4*) to perform *in vivo* FMO4 loss of function (LOF). We obtained KP mice that developed KP-driven lung adenocarcinoma with or without FMO4 LOF (*KP/shFmo4* or *KP/shNT*, respectively) (Extended Data Fig. 2a and 2b). 22 weeks after the lentiviral intratracheal delivery, we monitored tumor formation by computed tomography (CT) as we did before ^15^. Interestingly, the number of lung tumors was strongly decreased in *KP/shFmo4* mice compared with *KP/shNT* mice (total number of tumors = 14 vs 53 and mean number of tumors per mouse = 1.8 vs 5.9 in *KP/shFmo4* and *KP/shNT* mice, respectively) (Fig. 2a). At week 27, all *KP/shNT* mice had new detectable tumors, i.e., non-detected at week 22, but only 25% of *KP/shFmo4* mice did (total number of new tumors = 13 vs 3, respectively) (Fig. 2b). Moreover, the tumors already detected at week 22 grew more in *KP/shNT* than in *KP/shFmo4* mice (5.4-vs 3.6-fold at week 27) (Fig. 2c and 2d, Extended Data Fig. 2c). These data indicate that FMO4 LOF decreases both tumor generation and tumoral growth in this GEMM. And accordingly, the survival was increased in *KP/shFmo4* compared to *KP/shNT* mice (HR 4.584, 95% CI 1.531–13.719, *p* = 0.003) (Fig. 2e).

**Fig. 2.**
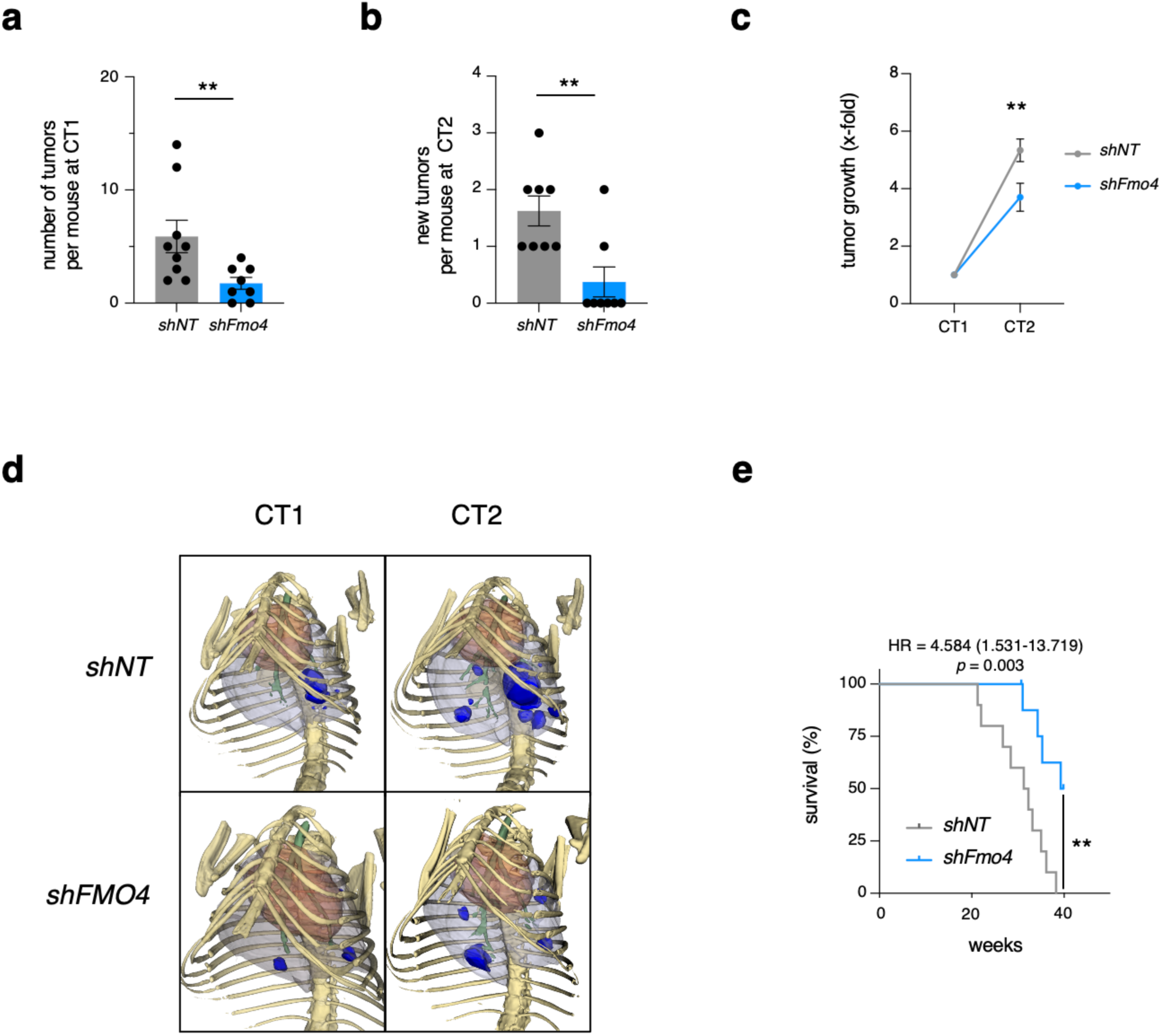
FMO4 is needed for oncogenic KRAS-driven lung adenocarcinoma development and growth. **a)** Number of tumor lesions per mouse identified by CT in control *shNT* (n = 9) and *shFMO4* (n = 8) KRAS^G12V^/p53^flox/flox^ mice at week 22 (CT1) after tumor induction. Data are the mean ± SEM; **p < 0.01, Mann-Whitney test. **b)** Number (mean ± SEM) of new tumor lesions identified by CT at week 27 (CT2) compared with the CT1 data in control *shNT* (n = 8) and *shFMO4* (n = 8) KRAS^G12V^/p53^flox/flox^ mice; **p < 0.01, Mann-Whitney test. **c)** Changes in volume of each individual tumor at CT2 relative to CT1 in control *shNT* (n = 32 tumors) and *shFMO4* (n = 11 tumors) KRAS^G12V^/p53^flox/flox^ mice. Results are presented as the mean ± SD; **p < 0.01, two-way ANOVA test. **d)** 3D reconstruction of examples of lungs containing CT-positive tumors from *shNT* and *shFMO4* KRAS^G12V^/p53^flox/flox^ mice. **e)** Kaplan-Meier survival curves for *shNT* (n = 10) and *shFMO4* (n = 8) KRAS^G12V^/p53^flox/flox^ mice. Statistical significance was calculated with the Mantel-Cox log-rank test: \*\**p* = 0.003 [HR = 4.584 (95% CI = 1.531–13.719)].

Altogether, our data demonstrate that FMO4 plays a major role in lung adenocarcinoma formation and homeostasis.

### Oxidative stress induces FMO4 expression through aryl carbon receptor (AHR) and FMO4 silencing increases reactive oxidative species (ROS) production

Oxidative stress regulation is crucial in aging ^29^ and ectopic expression of the FMO2 orthologue is sufficient to increase *C. elegans* life-span ^6^, furthermore same ectopic expression is sufficient to promote resistance against the oxidative stress inducer paraquat in this worm ^30^. As oxidative stress plays also a major role in cancer ^31^, we asked whether FMO4 could be implicated in oxidative stress protection.

First, we found that FMO4 was expressed at different levels in the tested human lung adenocarcinoma cell lines harboring different oncogenic driver mutations as well as in one non-transformed human lung cell line (Fig. 3a). Moreover, its expression level was positively correlated with ROS concentration (Fig. 3b). Then, to discern if it was cause or consequence, we induced ROS production in A549 cells (low FMO4 and ROS levels, Fig. 3b) by incubation with hydrogen peroxide (H_2_O_2_) producing a strong increase in FMO4 expression (Fig. 3c). Conversely, incubation of H1299 cells (high FMO4 and high ROS levels, Fig. 3b) with the ROS scavenger N-acetyl cysteine (NAC) led to a decrease of FMO4 expression (Fig. 3d). In both experiments, H_2_O_2_ and NAC concentrations were sublethal for these cell lines (Extended Data Fig. 3a). We conclude that FMO4 is positively regulated by oxidative stress.

**Fig. 3.**
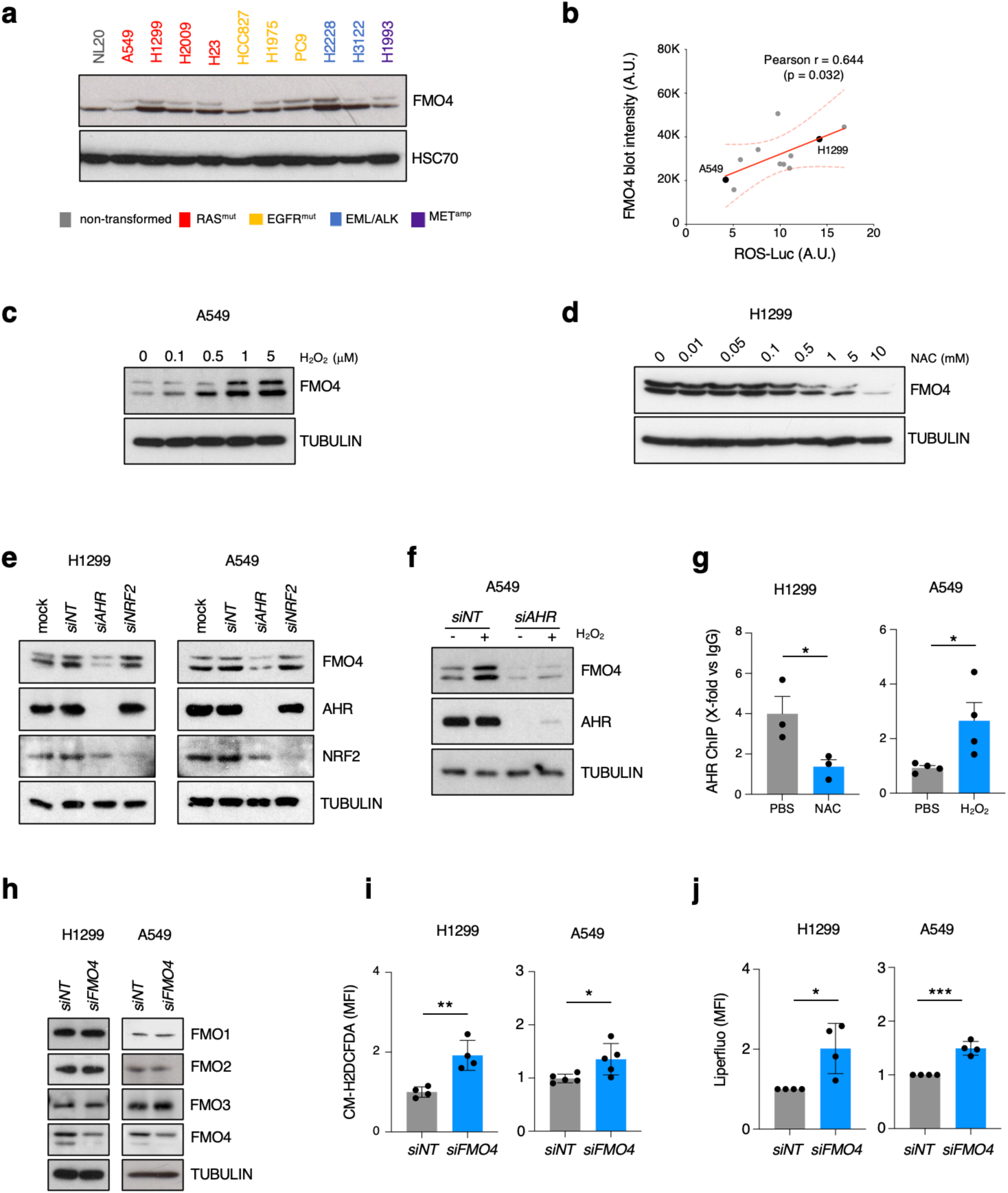
Oxidative stress induces FMO4 expression through AHR and FMO4 loss of function increases reactive oxidative species. **a)** Immunoblotting of the indicated proteins in different human lung cancer cell lines with different genetic alterations highlighted by different colors in the figure and in one non-transformed lung cell line (grey). **b)** Correlation between FMO4 expression and ROS levels in the same cell lines as in a). Pearson correlation coefficient = 0.644; *p* = 0.032. **c)** FMO4 expression in A549 cells incubated with the indicated concentrations of H_2_O_2_ for 24 h. **d)** FMO4 expression in H1299 cells incubated with the indicated concentrations of NAC for 24 h. **e)** Immunoblotting of the indicated proteins in H1299 cells (left panel) and A549 cells (right panel) transfected with a non-targeting siRNA (*siNT*), siRNAs against AHR (*siAHR*) or NRF2 (*siNRF2*), or mock-transfected (mock) for 48 h. **f)** Immunoblotting of the indicated proteins in A549 cells infected with siNT or siAHR for 48 h, and incubated or not with 5μM H_2_O_2_ for 24h. **g)** ChIP analysis of AHR binding to the *FMO4* promoter in H1299 cells incubated with 3mM NAC or PBS for 24h (n = 3) and in A549 cells incubated with 5μM H_2_O_2_ or PBS for 24h (n = 4). Data are presented as the mean ± SD; *p < 0.05, two-tailed unpaired Student’s *t*-test. **h)** Immunoblotting of the indicated proteins in H1299 (left panel) and A549 cells (right panel) transfected with a non-targeting siRNA (*siNT*) or siRNAs against FMO4 (*siFMO4*) for 48 h. **i)** Detection of reactive oxygen species (ROS) in cells transfected as in **h**, labelled with CM-H_2_DCFDA (10μM) for 1 h, and analyzed by FACS. Results are presented as the mean ± SD of normalized MFI; *p < 0.05, **p < 0.01, two-tailed unpaired Student’s *t*-test. **j)** Lipid peroxide quantification in cells transfected as in **h**, labelled with Liperfluo (10μM) for 1 h, and analyzed by FACS. Results are presented as the mean ± SD of normalized MFI; *p < 0.05, ***p < 0.001, two-tailed unpaired Student’s *t*-test.

To understand how ROS regulates FMO4 expression, we focused on the important regulator in response to oxidative stress in lung cancer: the nuclear factor erythroid 2– related factor 2 (NRF2) and its transcriptional inducer the aryl hydrocarbon receptor (AHR) ^9,32^. LOF of NRF2 in H1299 and A549 cells did not affect FMO4 levels (Fig. 3e), indicating that NRF2 is not required for FMO4 expression in this setting. In agreement with previous studies showing that NRF2 expression can be transcriptionally regulated by AHR ^32^, NRF2 levels were decreased upon AHR LOF, and of note, it also reduced FMO4 expression in both cell lines (Fig. 3e). Finally, we also exposed A549 cells to H_2_O_2_ or vehicle and treated them again with *siAHR* or *siNT*. Exposure to H_2_O_2_ increased FMO4 protein levels as before in *siNT* cells but not in *siAHR* cells (Fig. 3f). All our data demonstrate that AHR is implicated in controlling the oxidative stress-induced FMO4 expression.

To determine whether the transcription factor AHR directly binds to the *FMO4* promoter, we used the online http://alggen.lsi.upc.es/ tool to detect AHRE binding motifs in the *FMO4* promoter sequence and to design specific primers. Then, we performed chromatin immunoprecipitation (ChIP) experiments using an anti-AHR antibody and chromatin from H1299 cells (high ROS and FMO4 levels) incubated or not with NAC. Compared with the control immunoprecipitation (IgG), AHR immunoprecipitation resulted in enrichment of a *FMO4* promoter region containing one AHRE binding site in untreated cells (PBS), but not in cells incubated with NAC (Fig. 3g). Moreover, in A549 cells (low ROS and FMO4 levels), the same experiment showed little binding of AHR to the same AHRE-containing *FMO4* promoter region in untreated cells (PBS) and conversely, binding greatly increased upon incubation with H_2_O_2_ (Fig. 3g). Lastly, we assessed the correlation between *FMO4* and *AHR* mRNA levels in clinical lung adenocarcinoma samples using the GEPIA online tool ^33^, and found a positive correlation between both mRNAs (Extended Data Fig. 3b), suggesting that the AHR-FMO4 axis is acting also in human lung adenocarcinoma.

FMO4 was increased upon oxidative stress but its role in the response against this stress is unknown. Therefore, we used a pool of siRNAs against *FMO4* mRNA (*siFMO4*) in both H1299 and A549 cells. In *siFMO4* cells (both lines), the protein levels of FMO4 were effectively decreased, and interestingly neither FMO1, 2 nor 3 expression was affected, indicating that they did not compensate upon FMO4 LOF (Fig 3h). Importantly, oxidative stress was induced in both cell lines treated with *siFMO4* (Fig 3i), suggesting that FMO4 control oxidative stress. Then, and in order to test if this increased ROS levels could enhance the oxidation of cellular molecules, we also measured lipid peroxidation and observed that indeed Liperfluo was also increased in *siFMO4* treated cells compared to their *siNT* counterparts (Fig. 3j).

As a whole, our data reveal that FMO4 is regulated by oxidative stress through AHR and indicate that FMO4 counteract oxidative damage.

### FMO4 protects against ferroptosis *in vitro* and *in vivo*

Increased levels of lipid peroxidation can promote ferroptosis, a type of cell death implicated in tumor suppression ^11^ and with interesting perspectives for a therapeutic intervention in cancer ^10^.

To determine whether FMO4 was implicated in ferroptosis, we used a doxycycline-inducible lentiviral shRNA technology (TRIPZ), where shRNA induction upon incubation with doxycycline can be followed by monitoring the expression of a red fluorescent protein. First, we transduced H1299 and A549 cells with two different shRNAs against FMO4 (sh11 or/and sh12). Western blot analysis showed that in both cell lines, sh11+sh12 induced a stronger FMO4 decrease than any single shRNA, compared with the non-targeting shRNA (*shNT*) control (Extended Data Fig. 4a and Extended Data Fig. 4b). Therefore, we used the combination of both shRNAs for FMO4 LOF and called hereafter *shFMO4*. Then, we performed a clonogenic assay in both cell lines after transduction with *shFMO4* or *shNT*. In accordance with our *in vivo* LOF data presented in Figure 2, upon doxycycline addition to express the shRNAs, colony number and size were severely decreased in *shFMO4* H1299 and A549 cells (Fig. 4a and Extended Data Fig. 4c). Next, we used DAPI incorporation to test cell death, and interestingly the percentage of DAPI-positive cells was strongly increased in *shFMO4* condition in both cell lines upon doxycycline treatment, implying that FMO4 protect cells from dying (Fig. 4b and Extended Data Fig. 4d).

**Fig. 4.**
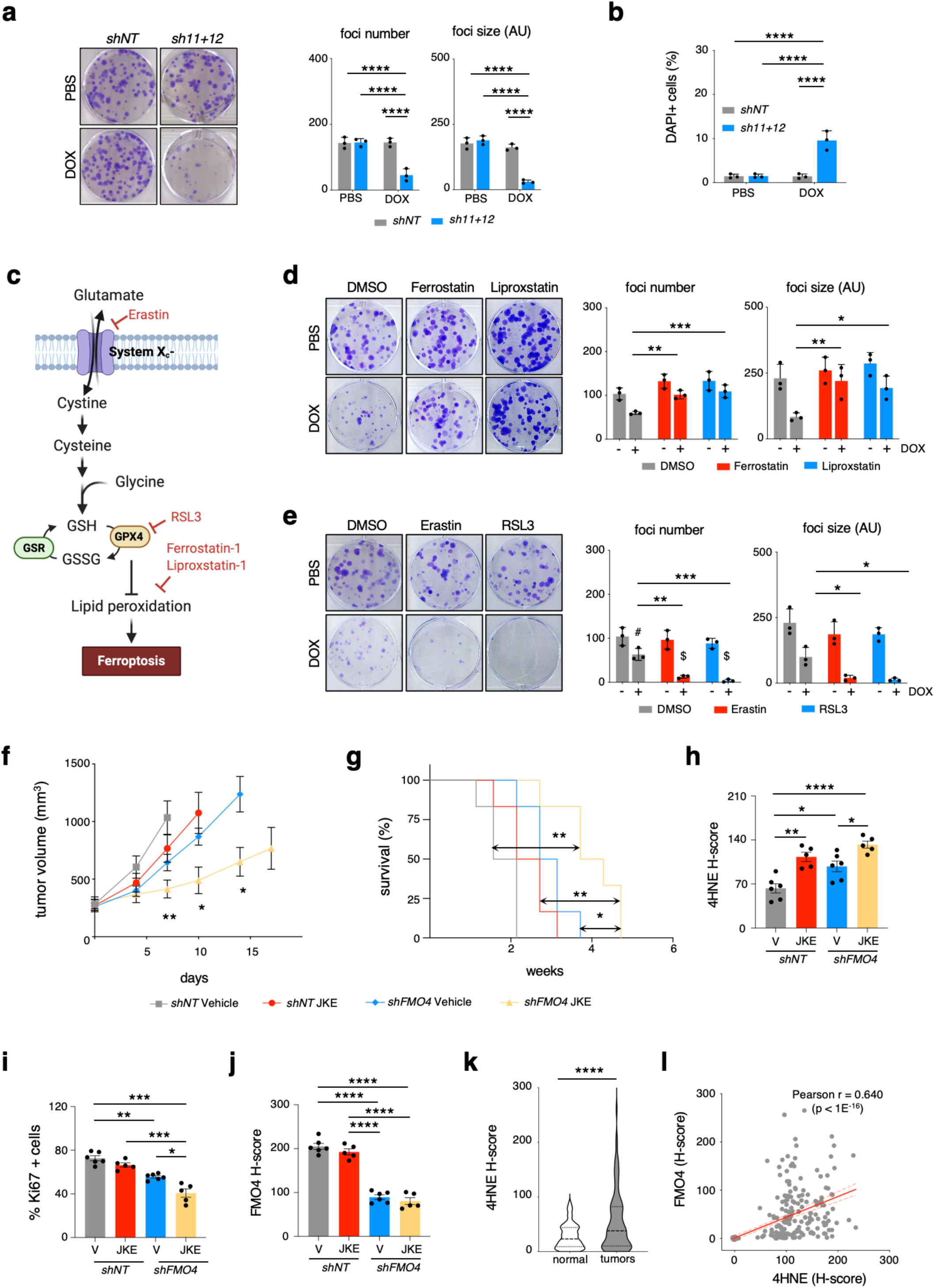
FMO4 protects against ferroptosis *in vitro* and *in vivo*. **a**) Representative images of clonogenic assays in control (*shNT*) and *sh11+12-H1299* cells. Cells were plated at low density and incubated with PBS or doxycycline (DOX; 1μg/ml) for 2 weeks. The number (left plot) and size (right plot) of foci in each condition are presented as mean ± SD (n = 3); \*\*\*\**p* < 0.0001, two-way ANOVA with Fisher’s LSD test. **b**) Quantification of the percentage of DAPI-positive cells in H1299 cells by flow cytometry (n = 3); \*\*\*\**p* < 0.0001, two-way ANOVA with Fisher’s LSD test. **c**) The cartoon depicts where Erastin and RSL3 (ferroptosis inducers) as well as Ferrostatin and Liproxstatin (ferroptosis inhibitors) act. GSR: glutathione reductase; GPX4: glutathione peroxidase 4. **d**) Representative images of clonogenic assays in H1299*-shFMO4* cells incubated with PBS or doxycycline (DOX; 1μg/ml) and with DMSO, Ferrostatin (5μm) or Liproxstatin (5μM) for 2 weeks. The number (left plot) and size (right plot) of foci in each condition are presented as mean ± SD (n = 3); * *p* < 0.05, ** *p* < 0.01, ***p < 0.001, two-way ANOVA followed by Sidák’s multiple comparison test. **e)** Representative images of clonogenic assays in H1299*-shFMO4* cells incubated with PBS or doxycycline (DOX; 1μg/ml) and with DMSO, Erastin (100nM) or RSL3 (20nM) for 2 weeks. The number (left plot) and size (right plot) of foci in each condition are presented as mean ± SD (n = 3); \**p* < 0.05, \*\**p* < 0.01, \*\*\**p* < 0.001, two-way ANOVA followed by Sidák’s multiple comparison test. **f)** Growth of H1299-*shNT* and -*shFMO4* tumor xenografts in mice fed with doxycycline-containing diet and treated with vehicle or JKE1674 (25mg/kg) daily 5 times/week (*shNT*-Vehicle n = 6, *shNT*-JKE n = 6, *shFMO4*-Vehicle n = 7, *shFMO4*-JKE n = 6). Mice were euthanized when tumors reached 1500 mm^3^ in volume. Statistical significance was calculated vs shFMO4-JKE condition with one-way ANOVA followed by the Bonferroni’s multiple test for comparison at day 7 and day 10; unpaired two-tailed Student’s *t* test for comparison at day 14.**g**) Kaplan-Meier survival curves of the mice described in **f**. Statistical significance was calculated with the Mantel-Cox log-rank test: \**p* < 0.05, \*\**p* < 0.01. **h)** Quantification of 4HNE staining (H-score) in tumors from the mice described in **f**. Results are presented as the mean ± SEM; #*p* < 0.1, \**p* < 0.05, \*\**p* < 0.01, \*\*\**p* < 0.001, two-way ANOVA followed by Sidak’s post hoc test. **i)** Quantification of the percentage of Ki67-positive cells in H1299-*shNT* and -*shFMO4* tumor xenografts from the mice described in **f**. Results are presented as the mean ± SEM; \**p* < 0.05, \*\**p* < 0.01, \*\*\**p* < 0.001, two-way ANOVA followed by Sidak’s post hoc test. **j)** Quantification of FMO4 staining intensity (H-score) in H1299-shNT and -shFMO4 tumor xenografts from the mice described in **f**. Results are the mean ± SEM; \*\*\*\**p* < 0.0001, two-way ANOVA followed by Sidak’s post hoc test. **k)** Violin plots showing the 4HNE H-score in normal lung (n = 244) and tumor (n = 185) biopsies. The median and upper/lower quartiles are depicted with dashed and dotted lines respectively; \*\*\*\**p* <0.0001, two-tailed unpaired Student’s *t*-test. **l)** Correlation between the FMO4 and 4-HNE H-scores (same TMA used in Fig. 3a). Pearson correlation coefficient = 0.64; \*\*\*\**p* < 0.0001.

We hypothesized that the above effects could be the consequence of ferroptosis, and to prove it, we used ferroptosis inhibitors (Ferrostatin and Liproxstatin) as well as inducers (Erastin and RSL3) (Fig. 4c)^11^. First, we performed cytotoxic assays in A549 and H1299 cells with these compounds to identify sublethal concentrations to be used for the experiments (Extended Data Fig. 4e). In agreement with our hypothesis, incubation of *shFMO4*-H1299 or *shFMO4*-A549 cells with any of the two ferroptosis inhibitors efficiently rescued the effect of FMO4 LOF on colony formation upon doxycycline exposition (Fig 4d and Extended Data Fig. 4f). Furthermore, incubation with the ferroptosis inducers strengthened the effect of FMO4 silencing (Fig 4e and Extended Data Fig. 4g), while once again, inhibitors alone at these concentrations had a very mild or no effect (Fig 4e and Extended Data Fig. 4g). Taken together, our *in vitro* data defined a protective role of FMO4 against ferroptosis.

To extend the link of FMO4 with ferroptosis to the *in vivo* setting, we selected JKE-1674, a highly specific GPX4 inhibitor with improved *in vivo* stability compared to previous compounds, such as RSL3 ^34^. First, we injected inducible *shFMO4*-H1299 or *shNT*-H1299 cells in the flank of immunodeficient NSG mice. When tumors reached a volume of ∼ 250mm^3^, we randomized *shFMO4*-H1299 and *shNT*-H1299 tumor bearing mice into two treatment groups (vehicle or JKE-1674) and switched to doxycycline-supplemented food to induce shRNA expression. We monitored tumor growth, as previously described ^16^, and stopped measurement in each arm when at least one tumor in that arm reached a volume of 1500mm^3^. This happened at day 7, 10, 14, and 17 in the vehicle-treated *shNT*-H1299 group, JKE-1674-treated *shNT*-H1299 group, vehicle-treated *shFMO4*-H1299 group, and JKE-1674-treated *shFMO4*-H1299 group, respectively (Fig 4f). Similarly, the mean survival time was 1.9, 2.4, 2.9, and 4 weeks in the vehicle-treated shNT-H1299 group, JKE-1674-treated shNT-H1299 group, vehicle-treated *shFMO4*-H1299 group, and JKE-1674-treated *shFMO4*-H1299 group, respectively (Fig 4g). Hence, FMO4 LOF potentiated the effect of the ferroptosis inducer JKE-1674 *in vivo*. Finally, at the used concentration, JKE-1674 did not affect mouse weight (Extended Data Fig. 4h).

Then, we analyzed for the expression of 4 hydroxynonenal (4-HNE), a surrogate ferroptosis marker that is generated by peroxidation of polyunsaturated fatty acids ^35^ and is frequently used in immunofluorescence/immunohistochemistry of mouse and human samples ^36,37^. In line with its role as ferroptosis inducer, 4-HNE was efficiently increased in tumors treated with JKE-1674 (Fig. 4h). We also measured proliferation, and found that the percentage of Ki67 cells progressively decreased in all groups compared with the vehicle-treated shNT-H1299 group (Fig. 4i). Lastly, we confirmed that FMO4 protein expression was reduced in *shFMO4*-H1299 tumors (Fig. 4j). Our *in vivo* data further confirm the protective role of FMO4 from ferroptosis, presenting this protein as an interesting target to sensibilize tumors against this type of cell death.

To test the clinical relevance of FMO4 role in ferroptosis, we quantified 4-HNE expression (H-score) in the same TMA used for FMO4 expression analysis above (Fig. 1d). Like FMO4, 4-HNE was strongly expressed in NSCLC samples compared with healthy lung tissue (Fig. 4k and Extended Data Fig. 4i) and more important, the FMO4 and 4-HNE H-scores showed a strong positive correlation (Fig. 4l).

Altogether, our data identified FMO4 as a novel protector against ferroptosis.

### FMO4 interacts with MAT2A and is required for the proper function of the methionine cycle

To understand how FMO4 protects against ferroptosis, we performed targeted metabolomics analysis ^38^ using *shNT*- and *shFMO4*-H1299 cells treated with doxycycline. We hypothesized that FMO4 LOF would promote the reduction of key molecules necessary for ferroptosis protection and hence we focused on metabolites that were decreased upon FMO4 silencing and analyzed them using the online tool https://www.metaboanalyst.ca/. We found two metabolic pathways with an FDR < 0.05: “cysteine and methionine metabolism” and “one carbon pool by folate” (Fig. 5a).

**Fig. 5.**
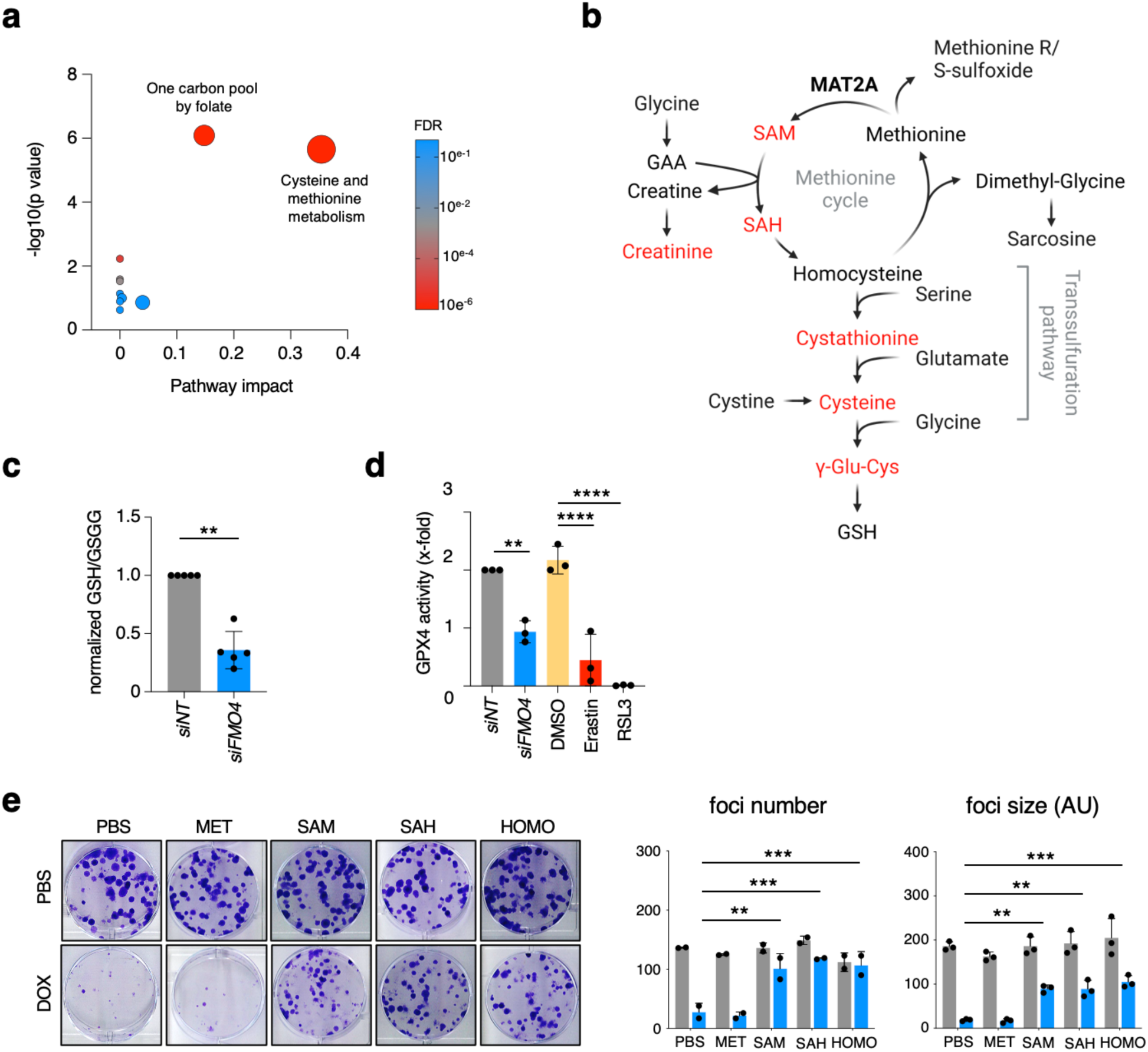
FMO4 interacts with MAT2A and is required for the proper function of the methionine cycle. **a)** Summary of pathway analyses determined by MetaboAnalyst 6.0 (https://www.metaboanalyst.ca), as visualized by bubble plots. Bubble sizes are proportional to the impact of each pathway, and bubble colors denote the degrees of significance, from highest (red) to lowest (blue). **b)** Schematic representation of the methionine and cysteine cycles. Metabolites downregulated upon FMO4 LOF (see Supp. Fig. 5a) are highlighted in red. **c**) Quantification of the GSH/GSSG ratio in H1299 cells transfected with a non-targeting siRNA (*siNT*) or siRNAs against FMO4 (*siFMO4*) for 48 h. Results are the mean ± SD (n=5); \*\**p* < 0.01, two-tailed unpaired Student’s *t*-test. **d)** Quantification of GPX4 activity in H1299 cells transfected as in **c** or incubated with DMSO, Erastin (2μM) or RSL3 (2μM) overnight. Results are the mean ± SD (n = 3); two-way ANOVA followed by Sidak’s multiple comparison test. **e**) Representative images of clonogenic assays in H1299-*shFMO4* cells incubated with PBS or doxycycline (DOX; 1μg/ml), and with methionine (MET, 500μM), S-adenosyl-methionine (SAM, 10μM), S-adenosyl-homocysteine (SAH, 50μM), or homocysteine (HOMO, 500μM) for 2 weeks. The number (left plot) and size (right plot) of foci in each condition are presented as mean ± SD (n = 2); \*\**p* < 0.01, \*\*\**p* < 0.001, two-way ANOVA followed by Sidák’s multiple comparison test.

Then, we also performed a quantitative proteomic analysis to identify proteins that interact with FMO4. We detected 36 proteins common between three independent experiments and with a Normalized Spectral Abundance Factor > 2 (for more details see the Methods section) (Extended Data Table 4). Notably, in this group of proteins appear methionine adenosyltransferase 2A (MAT2A), a crucial enzyme in the methionine cycle, that is present in both the “cysteine and methionine metabolism” and the “one carbon pool by folate” identified in our metabolomic analysis. The methionine cycle is implicated in cancer development and progression ^12^ and cooperates with the downstream transsulfuration pathway to generate cysteine that in turn produce reduced glutathione independently of the cystine uptake by system Xc^-^, the target of Erastin ^11^. Accordingly, the transsulfuration pathway inhibition sensitizes cells to Erastin ^39^ and hence, providing a putative explanation for the FMO4 protective role against ferroptosis. Figure 5b shows the methionine and transsulfuration pathways highlighting the metabolites found to be decreased in our targeted metabolomic analysis. Interestingly, these metabolites followed a pattern that could be explained by a defective MAT2A activity (Fig. 5b and Extended Data Fig. 5a). Conversely, cystine levels were not affected by FMO4 silencing (Extended Data Fig. 5a), indicating that the system Xc^−^ was not affected and hence further connecting the FMO4-mediated protection against ferroptosis with the methionine and transsulfuration pathways.

We also measured reduced glutathione and GPX4 activity, that should reduce upon decreased levels of FMO4 if our hypothesis is correct. Indeed, the reduced/oxidized glutathione ratio was decreased in H1299 cells *siFMO4* treated compared with controls (*siNT*) (Fig. 5c). Likewise, GPX4 activity was also strongly reduced in *siFMO4* treated H1299 cells, in a similar fashion to Erastin, while RSL3 totally abrogated the activity because it is a highly specific GPX4 inhibitor (Fig. 5d).

To further support our hypothesis, we incubated *shFMO4*-H1299 cells with the four products/substrates of the methionine cycle: methionine, SAM, SAH and homocysteine (Fig. 5b) at non-deleterious concentrations (Extended Data Fig. 5b). Remarkably, only methionine, the MAT2A substrate, could not rescue either foci number nor size in *shFMO4*-H1299 cells upon doxycycline exposure (Fig. 5d) while all the other 3 efficiently rescued FMO4 LOF as expected.

Taken together, these data revealed that FMO4 regulates MAT2A activity.

### FMO4 promotes MAT2A and MAT2B interaction

To understand how FMO4 regulates MAT2A activity, we used AlphaFold 3 ^40^ to model the different protein complexes, followed by molecular dynamics (MD) simulations to assess their stabilities and free energy of interaction calculations.

We started modelling the structures of the MAT2A homodimer (MAT2A_2_) and the MAT2A_2_/MAT2B complex because MAT2B binds to and stabilizes MAT2A ^41^ (Fig. 6a-b). The structures closely matched the corresponding X-ray crystallography structures ^42,43^, with Cα RMSD values of 0.348 Å for MAT2A_2_ and 0.614 Å for MAT2B (Extended Data Figs. 6a and 6b). The additional 25-aa N-terminal beta hairpin of MAT2B, modelled by AlphaFold 3 but absent in the X-ray structures, was aligned along the outside of one MAT2A monomer, and helped to stabilize their interaction (Extended Data Fig. 6b).

**Fig. 6.**
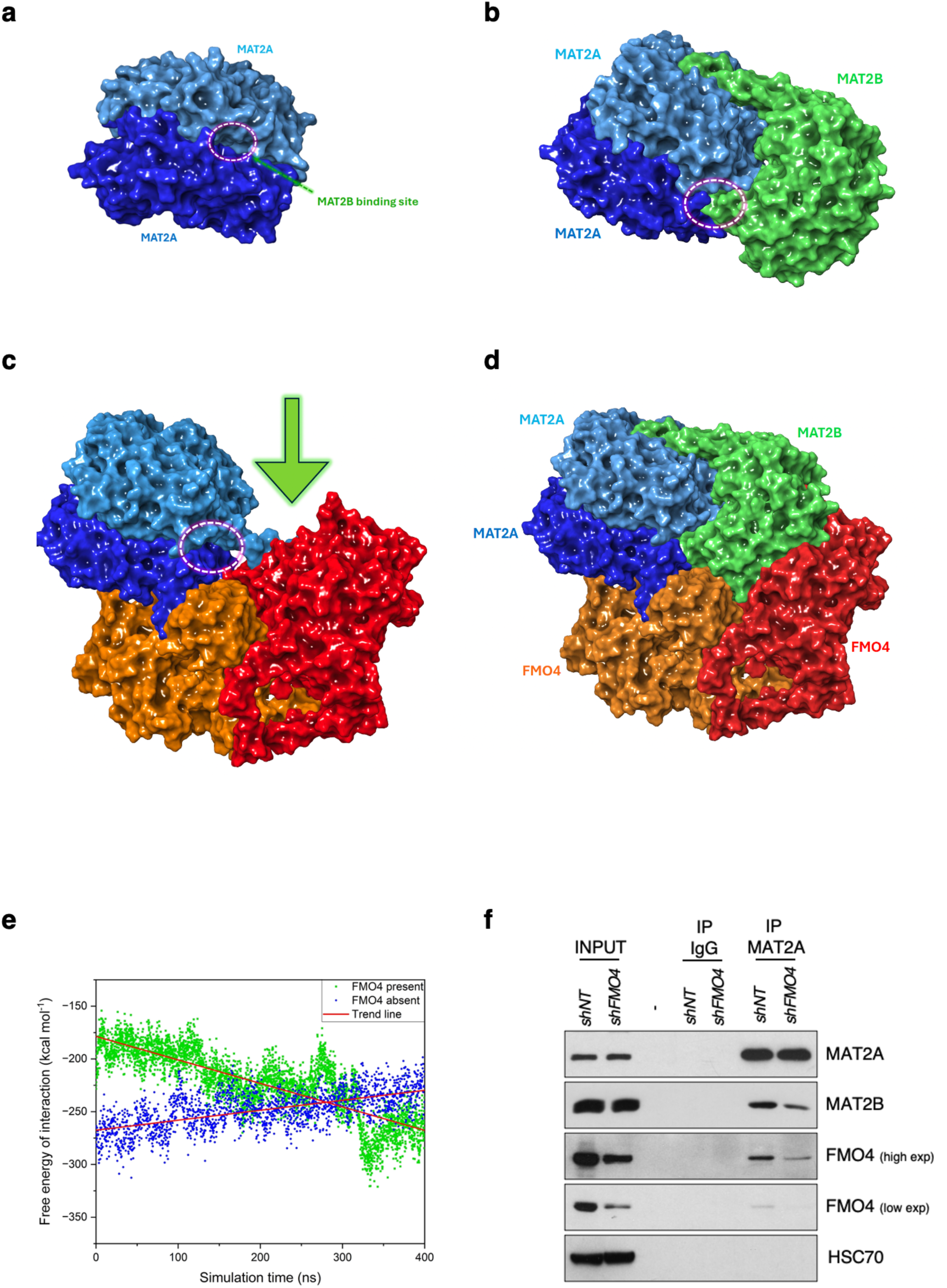
FMO4 promotes the interaction between MAT2A and MAT2B. Models of **a)** MAT2A_2_ (dark and light blue surfaces for each monomer) and **b)** MAT2A_2_/MAT2B complexes (green surface indicates MAT2B) generated by AlphfaFold3. MAT2A_2_ allosteric binding site that interacts with the C-terminal tail of MAT2B is highlighted by a pink dashed-line circle. **c)** The last snapshot of the 400 ns MD simulation for the MAT2A_2_-FMO4_2_ complex (the orange and red surfaces represent FMO4 monomers) where MAT2A_2_ was shifted with a tilting motion towards one of the FMO4 monomers, exposing the allosteric site (highlighted by a pink dashed-line circle), and generating a groove (green arrow) for accommodating MAT2B. **d)** The last snapshot of the 400 ns MD simulation of the MAT2A_2_-MAT2B-FMO4_2_ complex. **e)** The free energy of interaction (kcal mol^-1^) between MAT2A_2_ and MAT2B in the absence (blue) and presence (green) of FMO4_2_. The trend lines (red solid lines) indicate that FMO4_2_ stabilizes the MAT2A_2_-MAT2B complex. **f)** Immunoprecipitation with an anti-MAT2A antibody (IP MAT2A) or irrelevant IgG control (IP IgG) and then immunoblotting for indicated proteins of lysates from H1299-*shNT* and H1299-*shFMO4* cells incubated with doxycycline (DOX; 1μg/ml) for 72h h.

The AlphaFold 3 model quality was very satisfactory, according to the confidence scores (plDDT) and RMSD deviations from the corresponding X-ray structures. Nevertheless, we performed MD simulations to improve the model quality. The results clearly indicated that MAT2A_2_ and the MAT2A_2_/MAT2B complexes underwent a minor refinement during MD simulation by reaching a plateau in the corresponding Cα RMSD curves with values of 1.7 Å and 2.5 Å from the starting points (AlphaFold 3 models) for MAT2A_2_ and the MAT2A_2_/MAT2B complexes, respectively (Extended Data Figs. 6c and 6d). We next modelled the FMO4 homodimer (FMO4_2_) that showed a high structural similarity to the known dimeric X-ray crystal structures of yeast and bacterial FMOs (PDB ID 1VQW and 2XHV, respectively) ^44,45^. We also modeled the interaction of MAT2A_2_ and FMO4_2_, detected in our proteomic analysis (Extended Data Table 4). The initial AlphaFold 3 structure was highly symmetric and placed MAT2A_2_ with the side containing the allosteric cavity that interacts with the FMO4_2_ region furthest from the transmembrane helices. However, in the MD simulation, MAT2A_2_ shifted with a tilting motion towards one FMO4 monomer, thereby exposing the allosteric site and generating a groove to accommodate MAT2B (Fig. 6c and Extended Data Fig. 6e). Prompted by this intriguing finding, we modelled the full protein complex with FMO4_2_, MAT2A_2_ and MAT2B, that indeed fit perfectly in the groove (Fig. 6d). MAT2B C-terminal loop was inserted into the MAT2A_2_ allosteric pocket, and the N-terminal beta hairpin was aligned onto the outer MAT2A_2_. The 400 ns MD simulation of the full complex revealed a gradual adjustment and relaxation of the complex structure (Extended Data Fig. 6f).

Based on the simulation trajectories, we hypothesized that FMO4_2_ could facilitate MAT2B binding to MAT2A_2_, and calculated the free energy of interaction between MAT2A_2_ and MAT2B in the presence and absence of FMO4_2_. During the MD simulations, the free energy of interaction between MAT2A_2_ and MAT2B in the absence of FMO4_2_ tended to increase, *i.e.*, the interaction strength decreased with time. Conversely, in the presence of FMO4_2_ the free energy of interaction decreased with time, meaning that the strength of the MAT2A_2_-MAT2B interaction was enhanced (Fig. 6e). To support this hypothesis, we performed immunoprecipitation of endogenous MAT2A in H1299 cells and analyze the interaction with both FMO4 and MAT2B. First, we confirmed that MAT2A interacts with FMO4 as well as with MAT2B. Second, upon FMO4 LOF, FMO4 levels were decreased in the MAT2A immunoprecipitation, and remarkably, we also observed a strong decrease in MAT2B interaction with MAT2A, confirming the FMO4 role in stabilizing the MAT2A_2_-MAT2B complex (Fig. 6f).

Altogether, our data revealed that FMO4 plays a major role in the MAT2A_2_-MAT2B protein complex formation.

## DISCUSSION

State-of-the-art GEMMs that mimic human disease are crucial for advancing cancer knowledge ^46^. Thanks to the proteomic analysis of two widely used GEMMs of oncogenic EGFR- and KRAS-dependent lung adenocarcinoma ^15,16^, we identified FMO4, a protein with largely unknown functions in cancer, as highly expressed in EGFR-, KRAS- and EML4/ALK translocation dependent murine lung adenocarcinoma. These genetic alterations represent ∼50% of all driver oncogenic events described in human lung adenocarcinoma ^22^. FMO4 expression was four times higher in clinical lung cancer samples compared with normal lung, increasing the clinical relevance of our findings in mice. Moreover, FMO4 was amplified in 3-8% of patients, and its expression correlated with shorter 5-year overall survival. All these features point towards oncogenic functions for FMO4 in lung adenocarcinoma. In agreement, FMO4 loss of function in KP mice strongly reduced tumor burden and increased survival highlighting a crucial role in lung cancer for this protein. Following with this notion, before using the shRNA technology, we performed CRISPR in murine KRAS-driven lung adenocarcinoma cells to delete FMO4 and all clones died, further showing the importance of this protein in lung adenocarcinoma.

We also showed that FMO4 expression is increased by oxidative stress, which is regulated by several proteins, particularly the oxidative stress master regulator NRF2 ^9^. Our data indicated that NRF2 is not involved in controlling FMO4 expression, conversely AHR, a known NRF2 transcriptional inducer, binds to the *FMO4* promoter and activates FMO4 expression. Furthermore, FMO4 LOF strongly reduced colony formation, and this effect was eliminated by ferroptosis inducers. Lastly, we showed that ferroptosis inducers cooperate with FMO4 LOF *in vitro* but also *in vivo*, indicating that FMO4 is an interesting new target in lung cancer and we propose that its therapeutic inhibition could accelerate the use of ferroptosis inhibitors in the clinic. Taken together, our data suggest a feedback loop where oxidative stress increases FMO4 levels to control the damage associated with excessive oxidative stress. Furthermore, since both NRF2 and FMO4 are transcriptionally regulated by AHR, we hypothesize that FMO4 and NRF2 could work in parallel to control oxidative stress. Finally, our work is in accordance with previous studies showing AHR as a protector against ferroptosis ^47,48^.

Additional mechanistic insights linked FMO4 to MAT2A, a key enzyme in the methionine cycle that has a major role in cancer. Conversely, its role in ferroptosis is less clear. For instance, it was described that SAM, the production of which is catalyzed by MAT2A, is required to promote ferroptosis ^49^. Conversely, homocysteine reduction enhanced ferroptosis in lung cancer cells ^50^, and furthermore, the transsulfuration pathway, which is downstream the methionine cycle, can protect against ferroptosis ^39^. In both cases, authors proposed that that two different cysteine pools can be used by cells to generate reduced glutathione one coming from the Xc^−^ system, i.e., the target of Erastin, and another one from the methionine cycle. Our data fit better with the second set of studies because the strong decrease in clonogenic assay due to FMO4 loss of function can be rescued by incubation with SAM, SAH or homocysteine. Of note, we used SAM, SAH and homocysteine at concentrations non-deleterious to cells because SAM is very toxic at higher concentrations ^51^. This could also explain the disparities among different studies. Our work also fits with a previous study in *C. elegans* where the FMO2 orthologue in the worm was linked to the methionine cycle ^30^. Although the authors could not link the phenotype to any particular enzyme, together with our data suggest a conserved role for FMOs in the animal kingdom in controlling this key metabolic pathway that fits with the high degree of FMO protein family conservation during evolution ^3^.

We also modelled the interaction of FMO4 and MAT2A using AlphaFold 3, and observed that FMO4 forms dimers that interact with MAT2A dimers. This analysis revealed that the interaction with MAT2B is facilitated in FMO4 presence. Indeed, in cells with decreased FMO4 levels, MAT2B interaction with MAT2A was also decreased. Interestingly, a previous study showed that MAT2B increases MAT2A stability ^41^, particularly upon increased oxidative stress ^52^, and our data confirmed and extended these observations.

In summary, we identified FMO4, a largely unknown protein in cancer. Its wide presence in lung cancer developed by different oncogenes, as well as the strong effect promoted by its loss of function *in vivo* unveiled a new target in lung cancer and warrants the generation of specific inhibitors, either alone, in combination with ferroptosis inducers or with MAT2A inhibitors that are at the moment in clinical trials as AG-270 ^53^.

## MATERIAL AND METHODS

### Mice, genotyping and small imaging

The KRAS^G12D^;p53^flox/flox 28^, KRAS^G12V^;p53^flox/flox 54^, EGFR^L858R/T790M^/CCSP-rtTA ^16^ and EML4/ALK-driven lung adenocarcinoma ^18^ mouse strains were previously described. In the EGFR^L858R/T790M^/CCSP-rtTA strain, transgene expression was induced by feeding a doxycycline diet (1mg/kg) from SAFE. In KRAS^G12V^;p53^flox/flox^mice, tumors were induced by intratracheal delivery of adenovirus particles that express the Cre recombinase (3 x 10^8^ pfu) obtained from Iowa University, as previously described ^15^. In KRAS^G12D^;p53^flox/flox^mice, tumors were initiated by infection of lung cells with a lentiviral vector (pLV-U6:shRNA-mPGK:Cre) that delivers the Cre recombinase to activate oncogenic *Kras^G12D^* and delete *p53* ^28^. Ten-to fourteen-week-old mice were infected intratracheally with 10000 Cre-active lentiviral units. To induce EML/ALK tumor development, adenovirus particles that express Cas9 and the appropriate sgRNAs (1.5 x 10^8^ pfu), obtained from ViraQuest, were intratracheally delivered into CD1 mice, as previously described ^18^. Tumor appearance and progression were monitored by computed tomography using a nanoScan device (Mediso). The tumor area was analyzed with InterView™ FUSION (Mediso). The reconstructions of 3D images were made using 3D slicer software.

All animal procedures were performed according to protocols approved by the French national committee of animal care.

### Tandem Mass Tag (TMT) mass spectrometry

Murine lung samples were washed three times with ice-cold PBS and were prepared using the EasyPep Mini MS Sample Prep Kit (Thermo Fisher Scientific) according to the manufacturer’s instructions. Briefly, mouse lung tissue samples were transferred to Lysing Matrix M tubes (MP Biomedicals) with lysis buffer, and ground with an oscillation frequency of 6500 rpm for 1 min. Tissue extracts were clarified by centrifugation at 14000 rpm for 25 min before being immediately stored at −80°C until analysis. Protein concentration was measured using the Micro BCA Kit (Thermo Fisher Scientific). Total proteins (100 μg) were treated with Universal nuclease, and reduction and alkylation were achieved by incubation with the respective solutions at 95°C for 10 min. After addition of the trypsin/Lys-C protease mix, samples were incubated at 37°C for 3 h. Tryptic digestion was stopped before peptide clean-up and drying under vacuum. Dried peptides were labelled using the TMTpro 16-plex isobaric label reagents (Thermo Fisher Scientific) according to the manufacturer’s protocol. After quenching, samples were pooled and dried under vacuum. Before analysis, the peptide pool was fractionated into eight fractions using the High pH Reversed-Phase Peptide Fractionation Kit according to the manufacturer’s instruction (Thermo Fisher Scientific).

Peptides were analyzed online using a nano-flow HPLC (RSLC U3000, Thermo Fisher) coupled to a mass spectrometer equipped with a nano-electrospray source and the FAIMS technology (Exploris 480, Thermo Fisher). Peptides were separated following a gradient from 2 to 31% of buffer B (0.1% formic acid in 80% acetonitrile for 120 min at a flow rate of 300 nl/min, then 31 to 50% in 5 min, and finally 50 to 90% in 1 min. Tandem mass spectrometry analyses were performed in a data-dependent mode. For FAIMS, a gas flow of 4 l/min and CVs = −45 V and −60 V were used. Full scans (375– 1,500 m/z) were acquired in the Orbitrap mass analyzer with a resolution of 60,000 at 200 m/z. The most intense ions were sequentially isolated (isolation window of 0.7 m/z), fragmented by higher-energy collisional dissociation (normalized collision energy of 32) and detected in the Orbitrap analyzer (TurboTMT set to TMTpro reagent). Spectra were recorded with the Xcalibur 4.5 software (Thermo Fisher). MS data were deposited in the ProteomeXchange Consortium (http://www.ebi.ac.uk/pride/archive/) via the PRIDE partner repository ^55^ (data set identifier: PXD060578).

### Proteomic data analysis

Raw data were analyzed using MaxQuant (version 2.1.2.0) with standard settings ^56^. For TMT data, mass spectra were compared to *Mus musculus* entries from Uniprot (UP000000589 2020_06). A minimum peptide length of 7 amino acids and maximum of 2 missed cleavages were allowed for the tryptic digest. TMT16 was specified as a variable modification and the isobaric labels were bias-corrected according to the manufacturer’s certificate. Carbamidomethyl (C) was set as fixed modification and N-terminal protein acetylation and oxidation (M) were set as variable modifications. Reporter mass tolerance was set to 0.003 Da and identified peptides were filtered using their precursor intensity fraction (PIF). A false discovery rate (FDR) of 0.01 for proteins and peptides was required. Statistical analyses were performed with Perseus v2.0.7.0 ^57^. The reporter intensities were imported, log 2-transformed and normalized using median centring. Conditions were compared using the two-tailed Student’s *t*-test with Benjamini– Hochberg correction for multiple hypothesis testing (FDR of 0.05). Heatmaps and hierarchical clustering were performed using z-scores obtained from the original normalized values.

### Western blotting

For western blotting, cells were lysed in RIPA lysis buffer [1% Triton X-100, 0.5% sodium deoxycholate, 0.1% SDS, 150mM NaCl, EDTA 5mM, 50mM Tris, pH 7.4, 2mM Na_3_VO_4_, phosphatase inhibitors (Sigma) and complete protease inhibitor mix (Roche)]. Lysates were sonicated and cleared by centrifugation at 15.000g at 4 C° for 20 min. Protein concentration was determined using the BCA Protein Quantification Kit (Thermo Fisher). Equal amounts of protein lysates were separated by 9% SDS-PAGE and transferred to PVDF membranes (GE Healthcare). Membranes were blocked in 5% milk in saline tris buffer with 0.1% Tween 20 detergent (TBS-T) at room temperature (RT) for 1 h, followed by incubation with primary antibodies at 4 C° overnight. After three washes with TBS-T at RT of 5 min, secondary antibody in TBS-T/5% milk was added at RT for 1 h. Chemiluminescence signals were revealed with RevelBlotPlus (Ozyme) and autoradiography films (GE Healthcare). The following antibodies were used: anti-NRF2 (#12271) and anti-AhR (#83200) from Cell Signaling Technology; anti-FMO2 (#HPA028261), and -tubulin (#4026) from Sigma; anti-FMO3 (#orb228818) and -FMO4 (#orb97010) from Biorbyt; anti-FMO1 (sc-376924), anti-FMO5 (#sc-393732), anti-MAT2A (#sc-398917), anti-MAT2B (#sc-514069) and anti-HSC70 (sc-7298) from Santa Cruz Biotechnology. Secondary antibodies were horseradish peroxidase-linked anti-rabbit (#7074) and anti-mouse (#7076) IgG from Cell Signaling Technology. Tubulin and HSC70 were used as loading controls.

### RT-qPCR

Total RNA was isolated using the RNeasy Mini Kit (Quiagen). Complementary DNA (cDNA) was synthesized with SuperScriptIII (Invitrogen), and qPCR was performed using SYBR Green (Applied Biosystems) and the following primer pairs:

*Fmo1*: forward 5’-GCCAGTCTTTACAAGTCTGTGG-3’ and reverse 5’-TCCAGGAATAGAGAATTTGGCAC-3’.

*Fmo2*: forward 5’-CCCGGACTTCGCATCTTCTG-3’ and reverse 5’-GCGTCAAAGACAGTGCGTTG-3’.

*Fmo3*: forward 5’-ACTGGTGGTACACAAGGCAG-3’ and reverse 5’-ATGGTCCCATCCTCAAACACA-3’.

*Fmo4*: forward 5’-GATTGGAGCTGGCGTAAGTG-3’ and reverse 5’-TGTCAGCAAACTTCCACAGTC-3’.

*Actb*: forward 5’-ATGCTCTCCCTCACGCCATC-3’ and reverse 5’-CACGCACGATTTCCCTCTCA-3.

Real-time PCR was carried out on a LightCycler 480 II (Roche). Reactions were run in triplicate. Expression data were normalized to the geometric mean of the *Actb* housekeeping gene to control the expression level variability, and analyzed using the 2-DDCT method.

### Immunohistochemistry

Two previously described TMAs ^21,27^ and formalin-fixed paraffin-embedded (FFPE) H1299 cell xenograft sections were used for immunohistochemistry. Tissue sections (3μm thick) were de-paraffinized in xylene and ethanol, and rehydrated. For antigen retrieval, sections were incubated in citrate pH 6.0 antigen unmasking solution and heated at 100°C for 10 min and let cool down for 30 min at RT. To block endogenous peroxidase activity sections were incubated in 5% H_2_O_2_ at RT for 30 min, followed by protein blocking of non-specific epitopes with 5% normal horse serum. Slides were then incubated with primary antibodies against FMO4 (#HPA049100, Sigma), 4NHE (#NHE11-S, Alpha Diagnostic), Ki67 (#M720, DAKO) and cleaved caspase 3 (#9661S, Cell Signaling) at 4°C overnight. After washing with TBS-T, slides were incubated with the SignalStain Boost IHC Detection Reagent (Cell Signaling #8114) at RT for 30 min. After washing with TBS-T, signals were revealed using the DAB Kit (Vector, #SK-4100). Slides were counterstained with haematoxylin, dehydrated, and mounted. The staining percentage and intensity scores were calculated using the QuPath software ^58^. Briefly, the H-score^20^ represents a semiquantitative assessment of the staining intensity (using the signal in the adjacent normal tissue as the reference) and the percentage of positive cells. The range of possible score values vary from 0 to 300. The TMAs H-score was the mean of three different sections from the same sample, while for the FFPE H1299 xenograft H-score was the mean of three areas of the same sections.

The Kaplan-Meier method was used to estimate survival curves in function of the expression intensity. The Mantel-Cox test (Log-Rank model) was used to compare survival between different groups and to calculate the p-value, while the Cox regression analysis was used to assess the influence of variables on survival time and to calculate the Hazard Ratio (HR). Overall survival (OS) was defined as the time interval between the diagnosis and death.

### Plasmid constructs and lentiviral production for *Fmo4* knockdown *in vivo*

The shRNAs against *Fmo4* were designed using the BLOCK-iT RNA designer (Thermo Fisher) and the Board Institute shRNA design portal and adapted following the pSICOLIGOMAKER 1.5 program instructions (http://web.mit.edu/jacks-lab/protocols/pSico.html; created by A. Ventura, Memorial Sloan-Kettering Cancer Center) for optimal oligonucleotide design. *Fmo4* #sh1.2, # sh1.3, #shC1 and shNT (non-targeting) control oligonucleotides were annealed and ligated into a HpaI-XhoI digested pLV-U6:shRNA-mPGK:Cre dual-promoter lentiviral vector (kind gift by J. Faget) downstream of a U6 promoter.

#sh1.2: 5’-GGGACTATCTCAGAGAATT-3’

#sh 3.1: 5’-GCAATTCCTGAGCCCACAT-3’

#sh C1: 5’-GTGACAAATGTCTGCAAAGAA-3’

#sh NT: 5’-CCGCAGGTATGCACGCGT-3’

Lentiviral particles were produced by the Montpellier Vectorology Core facility. Briefly 293T cells were co-transfected with the lentiviral plasmid of interest (pLV-U6:shRNA-mPGK:Cre), pMD2G (VSV-G protein) and pCMVR8.74 (lentivirus packaging vector). Viral supernatants were harvested 48 h post-transfection, filtered and concentrated by ultracentrifugation. The viral titer was inferred from Cre activity, using immortalized mT/mG Cre-reporter mouse embryonic fibroblasts (MEFs), generously provided by J. Faget. MEFs (5×10^4^ cells per well) were plated in 24-well plates. The day after, cell number was determined in one well, then two-fold serial dilutions for each virus suspension, starting from 20 μl, were added in 1 ml of medium (DMEM supplemented with 10% foetal bovine serum (FBS), non-essential amino acids and pyruvate). 72 hours later, cells were harvested and the percentage of GFP^+^ cells was determined by flow cytometry (Cytoflex, Beckman Coulter). By representing the percentage of GFP^+^ cells on the Y-axis and the volume of the viral suspension on the X-axis and applying a linear regression line that intercepted 0 (Y=aX; r>0,98), the coefficient (a) of that curve multiplied by the number of cells determined on infection day yielded the number of active Cre particles per μl of concentrated virus.

### Cell culture, reagents and transfection

The, A549, H1299, H2009, H23, HCC827, H1975, H2228, H3122, H1993 and HEK293T cell lines were from the American Type Tissue Culture Collection (ATCC); the NL20 and PC9 cell lines were from Y. Yarden’s laboratory. The lung tumor-derived M8-GFP^rev^Tomato murine cell line was from J. Faget. NL20 cells were maintained in F12/Ham medium supplemented with glucose (2.7g/L), non-essential amino acids (0.1mM), insulin (0.005 mg/ml), EGF (10ng/ml), hydrocortisone (500ng/ml), glutamine (2mM), 10% FBS and antibiotics. HEK293T and M8-GFP^rev^Tomato cells were maintained in DMEM supplemented with 10% FBS and antibiotics. All the other lung cancer cell lines were maintained in RPMI 1640 supplemented with 10% FBS and antibiotics. Doxycycline hyclate (#D9891), L-methionine (#64319), *S*-(5’-Adenosyl)-L-methionine (#A7007), S-(5’-adenosyl)-L-homocysteine (#A9384), L-homocysteine (#69453), N-acetyl-L-cysteine (#A9165), and H_2_O_2_ (#H1009), from Sigma, were diluted in culture medium and used as indicated. The non-targeting siRNA control (siNT) and the siRNAs against AHR (siAHR) and NRF2 (siNRF2) (purchased as smart pools from Dharmacon) were transfected at 20nM with Dharmafect2 reagent following the manufacturer’s instructions. Ferrostatin-1 (#SE-S7243), Liproxstatin-1 (#SE-S7699), Erastin (#SE-S7242) and RSL3 (#SE-S8155) were obtained from Selleckchem.

### Oxidative stress analysis in cell lines

ROS production was measured using the ROS-Glo™ H_2_O_2_ Assay kit (Promega) according to the manufacturer’s instruction. 15-20 x10^4^ cells were plated in opaque white 96-well plates, and 20µl of H_2_O_2_ substrate solution was added to the cells, to a final volume of 100 µl. Cells were incubated at 37°C in a 5% CO_2_ incubator for 1 h before addition of 100 µl of ROS-Glo detection solution to each well. After incubation at RT for 20 min, luminescence was measured using a EnSpire Alpha® luminometer (PerkinElmer).

### Chromatin immunoprecipitation (ChIP)

Chromatin was prepared as described previously ^59^. The ChIP-Adem-Kit and ChIP DNA Prep Adem-Kit (Ademtech) were used for ChIP and DNA purification, respectively, on an AutoMag robot, according to the manufacturer’s instructions. The anti-AHR antibody was from Cell Signaling (#83200) and the IgG rabbit control from Millipore. The immunoprecipitated DNA was analyzed by PCR using the following primers targeting the *FMO4* promoter: forward 5’-GCCAATCCAACAGCTGTATTCT-3’ and reverse 5’-GCCCTCAGTTTAAAACAAAAGC-3’.

### Flow cytometry analysis of oxidative stress or lipid oxidation

H1299 and A549 cells were transfected with siRNAs targeting FMO4 (siFMO4) or non-targeting (siNT) control for 48 h. To measure oxidative stress, whole cell suspensions were incubated with 10 μM CM-H_2_DCFDA (#C6827 ThermoFisher) for 1 h, according to the manufacturer’s instructions. DAPI (1μg/ml) was added just before the analysis. To measure lipid oxidation, cell suspensions were incubated with 5 μM Liperfluo (#L248 Dojindo) for 30 min, according to the manufacturer’s instructions. Cells were analyzed by flow cytometry (Cytoflex, Beckman Coulter) and the CM-H_2_DCFDA or Liperfluo mean fluorescence intensity (MFI) signals in viable cells were calculated using the FlowJo software.

### Clonogenic assay

Cells were seeded in 6-well plates in duplicate or triplicate at a density of 200-400 cells/well (according to cell type) in 2 ml of medium containing 10% FBS and antibiotics. After 24 h, medium was replaced with fresh medium containing 2% FBS in the presence or absence of doxycycline and/or the indicated compounds, and cultured at 37°C in a humidified atmosphere containing 95% air and 5% CO_2_ for 2 weeks. Cells were fixed with methanol and stained with a solution containing 0.5% crystal violet and 25% methanol for 15 min, followed by three rinses with water to remove excess dye. Colony number and size were analyzed using ImageJ.

### Viability assay

For sulforhodamine B (SRB), cells were plated in 96-well plates in quadruplicate or pentaplicate at the desired density and treated as indicated. At the end of the treatment period, culture medium was removed and cells were fixed with cold 10% trichloroacetic acid (Sigma) at 4°C for 1 h. Then, cells were washed five times with water, and plates were left to dry. Plates were stained with 200 µl 0.1% SRB dissolved in 1% acetic acid for at least 15 min, followed by four washes with 1% acetic acid to remove unbound stain. Plates were left to dry at RT and bound dye molecules were solubilized with 200µl of 10mM tris(hydroxymethyl)aminomethane (TRIS base) solution at pH 10.5. Optical density was measured at 540nm absorbance (PherastarFS, BMG Labtech).

For the DAPI exclusion, dead and dying cells amount was assessed with 4’,6-diamidino-2-phenylindole (DAPI from Sigma) exclusion staining. Cells were analyzed by flow cytometry (Cytoflex, Beckman Coulter) and the percentage of dead/dying cells was determined with the FlowJo software.

### Generation of cell lines that stably express pTRIPZ –FMO4 shRNA

The pTRIPZ doxycycline-inducible non-targeting shRNA (shNT) and shRNAs against FMO4 [clone V3THS_385089 (#89), clone V3THS_385090 (#90), clone V3THS_385093 (#93), clone V3THS_385094 (#94), clone V2THS_113911 (#11), clone V2THS_113912 (#12)] were purchased from Dharmacon. To generate stable cell lines, HEK293T cells were transfected with the pTRIPZ lentivirus vectors, psPAX2, and pVSV-G using lipofectamine 2000 (Life Technology) for 6-8 h. Medium was changed to DMEM with 10% FBS. Viruses were collected 48 and 72 h after transfection. shNT, sh11, and sh12 lentiviral particles were used immediately to transduce A549 and H1299 cells, or stored at −80 C°. Cells were infected in the presence of polybrene (5-10μg/ml), and selected by incubation with puromycin (2μg/ml) (Sigma) for up to 1 week. Cells were then incubated with 0.5μg/ml doxycycline overnight, and RFP^+^ cells were sorted using an ARIAIIIU (Becton Dickinson) cell sorter. Sorted cells were maintained in culture in doxycycline-free medium. For functional assays, the tetracycline responsive element (TRE) was induced by incubation with 1μg/ml (H1299 cells) and 1.5μg/ml (A549 cells) doxycycline as indicated.

### Mouse xenografts

2 x 10^6^ H1299 cells (infected with shNT or shFMO4) were injected subcutaneously in the flank of 6-10-week-old, male and female NSG mice (Charles River). When tumors reached the volume of ∼250 mm^3^, mice were randomized in four groups and fed ad-libitum with doxycycline-supplemented (1 mg/kg) food pellets (SAFE), and received the indicated drug treatments. Tumor volumes were calculated with a caliper and according to the formula V = (D x d^2^)/2, where D and d are the major and minor perpendicular tumor diameters, respectively. Mice were killed when tumors reached the volume of 1500 mm^3^. JKE1674 (#T37314; TargetMol) was resuspended in 0.5% (w/v) methylcellulose/0.2% Tween-80 (w/v) (vehicle) and administered (25 mg/kg/day) by oral gavage 5 days per week.

### Metabolomic analysis

For the metabolomic analysis, control (shNT) and sh11+12-infected H1299 cells were plated in 6-well plates and cultured in presence of doxycycline (1μg/ml) for 72 h. Cells were harvested by solubilization in 750 µL of water/methanol: 30/70 with 0.1% formic acid and 5 µL of 100 µM 2-morpholinoethansulfonic acid (Cat. # 341-01622; Dojindo, Tokyo, Japan) as internal standard. Glass beads were added in each tube and samples were vortexed vigorously for 1 min, homogenized with a Tissue Homogenizer Precellus 24, and centrifuged at 15 000 g at 4 °C for 10 min. Supernatants were loaded on a Captiva EMR plate (Agilent, Cat.# 5190-1001) assembled on the vacuum manifold (Agilent, Cat.# A796) together with the deep well collection plate (Agilent, Cat.# A696001000). The flow-through was collected and dried using a Speedvac concentrator. Samples were resuspended in 100µL of water, injected onto the liquid chromatography-mass spectrometry (LC-MS) system that consisted of ultra-high performance LC instrument (Agilent, 1200 Infinity II Biocompatible) coupled to a triple quadrupole MS instrument (Agilent, 6495C), and analysed with the MRM acquisition method. For more details on metabolomic analysis refer to (PMID: 37590149).

### Co-immunoprecipitation

For the identification of proteins interacting with FMO4, HEK293 cells were transfected with the pCMV6-FMO4-Myc-DDK plasmid (Origene #RC201181) or pCMV6-empty vector using Lipofectamine 2000 (Thermo Fisher). After 48 h, co-immunoprecipitations were performed according to the published STAR protocol ^60^. A magnetic bead-conjugated anti-DDK antibody (Origene #TA150042S) was use for the immunoprecipitation. Elution was performed using 0.5mM ETDA and 0.5M NH_4_OH (pH11.0) basic buffer.

For detecting MAT2A complexes, H1299 cells were incubated as indicated and lysates were prepared using 20mM Tris (pH7.5), 0.15 M NaCl, 0.2mM EDTA, 0.5% NP40, 0.5% Triton X-100, and 5% glycerol. Then, complexes were immunoprecipitated with the anti-MAT2A (#sc-398917) antibody and protein A/G plus (#SC-2003) agarose beads at 4°C overnight. An irrelevant antibody (IgG, #sc-3878) was used as control. Complexes were washed in HNTG buffer [20 mM Hepes (pH 7.5), 0.15 M NaCl, 0.1% Triton X-100, and 10% glycerol], resuspended in Laemmli buffer, and boiled for 10 min.

### Mass spectrometry of proteins interacting with FMO4

For protein identification, samples were separated on a 10% Mini-Protean TGX Tris-Glycine Precast Gel (Biorad). After a short migration and staining with Coomassie, a single band was excised. Each gel slice was washed, reduced with 10 mM DTT, alkylated with 55 mM iodoacetamide, and subjected to in-gel trypsin digestion overnight (Trypsin Protease, MS Grade Pierce, Thermo Fisher Scientific). The extracted tryptic peptides were cleaned up using OMIX C18 100 µl pipette tips (Agilent), lyophilized and reconstituted for the LC-MS/MS analysis. The resulting peptides were analyzed online by nano-flow HPLC (RSLC U3000, Thermo Fisher) coupled to MS-nano-electrospray ionization using a Qexactive HFX mass spectrometer (Thermo Fisher Scientific) coupled to a nano-LC system (Thermo Fisher Scientific, U3000-RSLC). Sample desalting and preconcentration were performed online on a Pepmap® precolumn (0.3 × 10mm; Fisher Scientific, 164568). A gradient consisting of 0% to 40% B in A (A: 0.1% formic acid [Fisher Scientific, A117], 6% acetonitrile [Fisher Scientific, A955], in H_2_O [Fisher Scientific, W6], and B: 0.1% formic acid in 80% acetonitrile) for 120 min (300nl/min) was used to elute peptides from the capillary reverse-phase column (0.075 × 250mm, Pepmap®, Fisher Scientific, 164941). Data were acquired using the Xcalibur software (version 4.0). A cycle of one full-scan mass spectrum (375–1,500 m/z) at a resolution of 60000 (at 200 m/z) followed by 12 data-dependent MS/MS spectra (at a resolution of 30000, isolation window 1.2 m/z) was repeated continuously throughout the nano-LC separation. MS data were deposited in the ProteomeXchange Consortium (http://www.ebi.ac.uk/pride/archive/) via the PRIDE partner repository ^55^, with the data set identifier PXD060586.

### Computational modeling of protein complex

The artificial intelligence-based structure modeling tool AlphfaFold3 (AF3) ^40^ was used to generate the initial models of FMO4 and its complexes. The amino acid sequences of human MAT2A, MAT2B and FMO4 were retrieved from UniProt (www.uniprot.org), with the entry codes P31153, Q9NZL9, and P31512, respectively.

For benchmarking, the structures of the MAT2A homodimer (MAT2A_2_) and the complex of MAT2A_2_ with MAT2B was modeled. The obtained complexes were superposed on the corresponding X-ray crystal structures, PDB IDs 5A1I (MAT2A_2_; 1.09 Å resolution) ^42^ and 4NDN (MAT2A_2_/MAT2B; 2.34 Å resolution) ^43^. The root mean square deviations (RMSD) of Ca backbone atoms between the AF3 models and the crystal structures were 0.348 in MAT2A_2_, and 0.614 Å for MAT2B in the larger complex, giving confidence in the method. Including post-translational modifications or co-factors, as listed in the UniProt entries, did not affect the structures.

The major difference in the various AF3 models of MAT2A_2_ was the orientation of the first N-terminal 12 residues (not resolved by X-ray crystallography) that formed either a well-structured a-helix extending out from the protein, or a less ordered chain aligned along the side of the protein. In the larger complex, the C-terminal tail of MAT2B extended down in the allosteric cavity between the two MAT2A monomers, perfectly aligned with the crystal structure. In addition, in the AF3 models, the 25 residues of MAT2B N-terminus, also not resolved in the crystal structures, formed a beta hairpin motif aligned as a ‘tongue’ along the outside of one of the MAT2A monomers, helping to keep MAT2B in position (Fig. 6b).

For the FMO4 homodimer (FMO4_2_) modeled with AF3, the per-residue confidence score (pLDDT) was generally high (mostly > 90), except for the specific orientation of the C-terminal transmembrane a-helix because the model was generated in the absence of the lipid bilayer.

The molecular dynamics (MD) simulations of MAT2A_2_ and the MAT2A_2_/MAT2B, MAT2A_2_/FMO4_2_ and MAT2A_2_/MAT2B/FMO4_2_ complexes were conducted using the Desmond module of Schrödinger 2024-1 (www.shrodinger.com) ^61^. Before the MD simulations, the structures retrieved from AF3 were carefully prepared using the Schrödinger protein preparation wizard ^62^ to correctly assign the bonds and bond orders and add missing hydrogens. The hydrogen bonding network was optimized by adjusting the protonation states of Asp, Glu and tautomeric states of His to match a pH of 7.0 ± 2 ^63^. Lastly, the structures underwent geometry refinement using the OPLS4 force field ^64^ in restrained structural minimization with a maximum all-atom RMSD of 0.3 Å. All MD simulations were performed for 400 ns in NPT ensemble employing the OPLS4 force field ^64^. Water molecules were modeled with the TIP3P force field ^65^. Periodic boundary conditions were enforced in all directions, and a 10 Å water buffer surrounded the protein within a cubic simulation box. The system net charge was neutralized by adding the appropriate number of counterions (Cl^−^/Na^+^), and the salt concentration was maintained at 150 mM to simulate physiological conditions. Temperature (300 K) and pressure (1 atm) were regulated using the Nose–Hoover thermostat ^66^ with a 1 ps relaxation time and the Martyna–Tobias–Klein barostat ^67^ with a 2 ps relaxation time and isotropic coupling, respectively. The short-range van der Waals and electrostatic interactions were computed using a 12–6 Lennard-Jones potential and Coulomb’s law, respectively, within a 10 Å cut-off radius. Long-range electrostatic forces beyond the cut-off radius were calculated using the smooth particle mesh Ewald (PME) method. The default initial minimization and relaxation protocol included the following steps:

a. NVT Brownian dynamics with solute heavy atom restraints at 10 K for 100 ps,
b. NVT simulation at 10 K with solute heavy atom restraints for 12 ps,
c. NPT MD simulation at 10 K with solute heavy atom restraints for 12 ps,
d. NPT MD simulation at 300 K with solute heavy atom restraints for 12 ps, and
e. NPT MD simulation at 300 K without restraints for 24 ps. Following minimization and relaxation, each system underwent a 400 ns production simulation.

The free binding energy of interactions for the MAT2A_2_/MAT2B complexes in the presence and absence of FMO4_2_ were calculated using the Molecular Mechanics Generalized Born Surface Area (MM-GBSA) method ^68^, implemented in the Schrödinger 2024-1 package. To improve the result reliability and accuracy, MM-GBSA calculations were conducted on MD trajectories at 1 ns interval, resulting in 400 snapshots per trajectory. The same force field and cut-off distances used in the MD simulations were maintained throughout the calculations.

The coordinate files of the different predicted complex structures and the MD simulation trajectories are freely available at Zenodo.org, through DOI 10.5281/zenodo.14840279.

### Statistical analysis

Unless otherwise specified, data are presented as means ± SEM. Two-way analysis of variance (ANOVA) followed by the Fisher’s LSD test was performed to compare tumor growth in Figure 2c and to compare the clonogenic and apoptosis assay results in Figure 4a, 4b and in extended data Fig.4c, 4d. One-way ANOVA followed by the Tukey’s post hoc test was performed to analyze the data in extended data Fig. 1c. One-way ANOVA followed by the Sidak’s post hoc test was used to analyze the findings shown in extended data Fig. 2b. In Figures 4d, 4e, 4h, 4i, 4j, 5d, 5e and in extended data Figure 4f, 4g, data were analyzed by 2-way ANOVA followed by the Sidak’s post hoc test. In Figure. 4f, data were analyzed by one-way ANOVA followed by the Bonferroni’s multiple test for comparison at day 7 and day 10; unpaired two-tailed Student’s *t* test for comparison at day 14. In extended data Figure 1a, data were analyzed by 2-way ANOVA followed by the Dunnett’s multiple comparison test. In Figures 1e, 2e, 4g and in extended data Figure 1f, the Kaplan-Meier survival curves were analysed with the Log-rank (Mantel-Cox) test. Hazard ratios were calculated using the log-rank test. The correlation between groups in Figures 3b 4i, and extended data Figure 3b was calculated using the Pearson *r* correlation coefficient. In Figures 1d, 3g, 3i, 3j, 4k, 5c and in extended data Figures 1e, 5a, data were analyzed with the unpaired two-tailed Student’s *t* test. In Figures 2a, 2b, data were analyzed with the Mann-Whitney test.

Samples (patients, cells, mice) were allocated to their experimental group according to their predetermined type. All statistical analyses were performed using the Prism 10 GraphPad software, except for extended data Figure 3b. Investigators were blinded to the experimental groups in the analysis of the data presented in Figure 2a, 2b, 2c, 4h, 4i, 4j and in extended data Figure 2b, 2c. The investigators were not blinded in the remaining analyses. *p* ≤ 0.05 was considered significant. # *p* < 0.1, * *p* ≤ 0.05, ** *p* ≤ 0.01, *** *p* ≤ 0.001, **** *p* ≤ 0.0001.

## Supporting information

ExtendedFigures

## ACKNOWLEDGMENTS

We thank Daniel Herranz, Raúl Durán, José Francisco Rodriguez, Manuel Collado, Alejo Efeyán, Xavier Guillory, Eric Chevet, Alexander Djiane, Patrick Joundin and Laurent Le Cam for helpful discussion and/or critical reading of the manuscript. Elisabetta Andermarcher professionally edited the manuscript. We thank Mariano Barbacid, Andrea Ventura, and Kwok Kin Wong for providing us the KRAS^G12V^/p53^flox/flox^, EML4/ALK and EGFR GEMMs respectively.

We acknowledge the “Réseau d’Histologie Expérimentale de Montpellier” - RHEM facility supported by SIRIC Montpellier Cancer Grant INCa_Inserm_DGOS_12553, the European regional development foundation and the Occitanie region (FEDER-FSE 2014-2020 Languedoc Roussillon), REACT-EU (Recovery Assistance for Cohesion and the Territories of Europe), IBiSA and Ligue contre le cancer for processing our animal tissues, histology technics and expertise. Mass spectrometry experiments were carried out using the facilities of the Montpellier Proteomics Platform (PPM-PP2I, BioCampus Montpellier), and were partially supported by the French National Research Agency ProFI fundings (Proteomics French Infrastructure, ANR-24-INBS-0015). The Orbitrap Exploris 480 mass spectrometer used for proteomics analysis was co-financed by the European Regional Development Fund (ERDF) and the Occitanie region (SAPHIR project). French Ministry of Higher Education and Research for supporting the Platform for Translational Oncometabolomics (PLATON) through a CPER grant (IBDLR). We acknowledge the imaging facility MRI, member of the France-BioImaging national infrastructure supported by the French National Research Agency (ANR-10-INBS-04, «Investments for the future»). We thank the vector facility, PVM, Biocampus Montpellier, CNRS UMS3426 for the lentivirus production. We thank the IRCM animal facility members, for their outstanding work that we extend to the small imaging platform (IPAM).

M.M. was supported by contracts from *Fondation ARC*, *Institut National du Cancer* (INCa_9257) and Occitanie Region/FEDER funds (LR0014155) and by INCa-Cancéropôle GSO. Work in A.M.’s lab is supported by the *European Commission* (EIC-TRANSITIONOPEN-ID101058055), the International Research Program (IRP) from *INSERM* and the *Institut National du Cancer* (INCA_16696). The funders had no role in the study design, data collection and analysis, decision to publish, or preparation of the manuscript.

