## ExtendedFigures for "FMO4 drives lung adenocarcinoma by stabilizing the MAT2A/MAT2B complex and hindering ferroptosis"

### EXTENDED DATA FIGURES AND LEGENDS

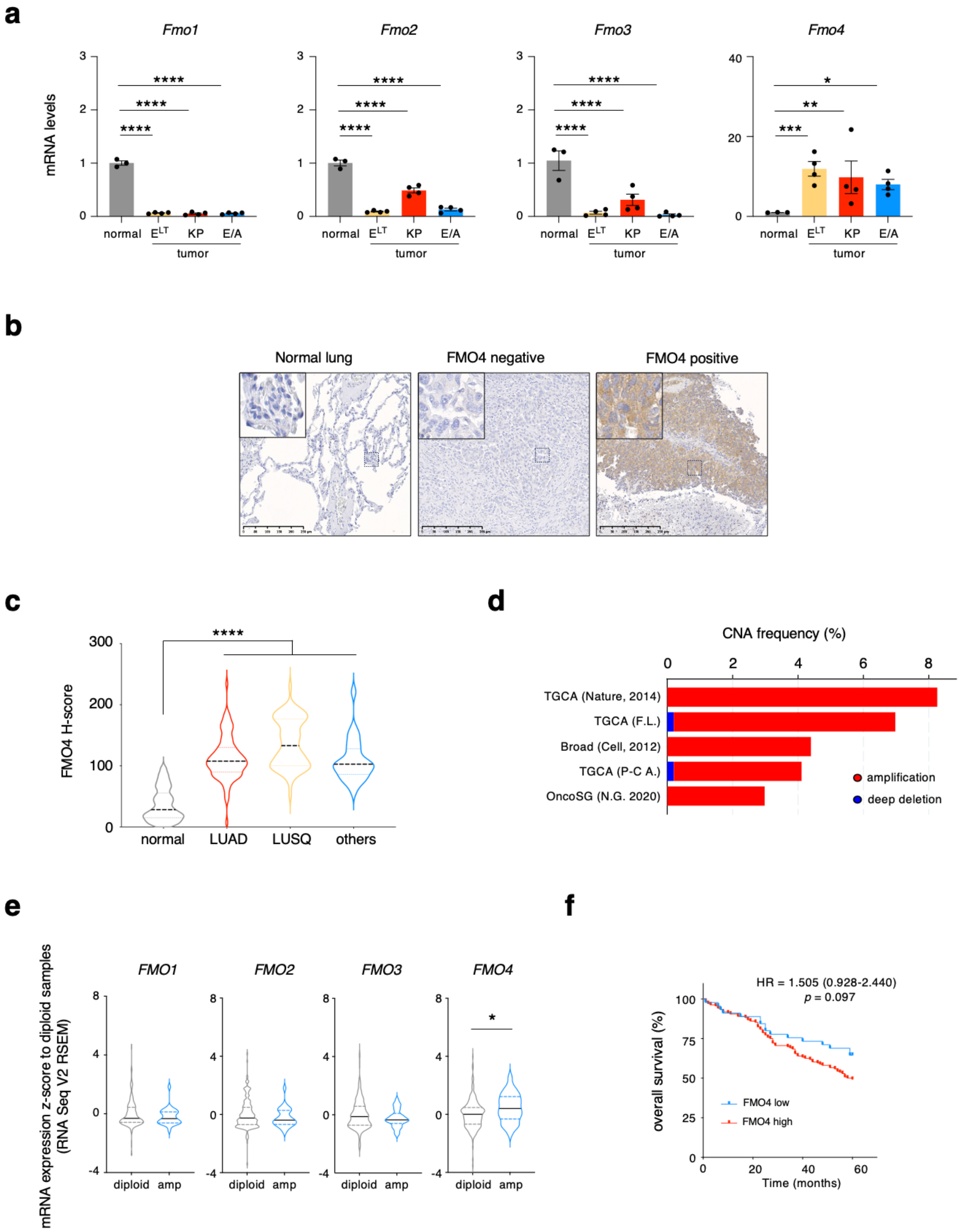


Extended Data Fig. 1, Bracquemond et al.

**Extended Fig. 1. FMO4 is highly expressed in mouse and in human lung cancer.**

**a)** *Fmo1*, *Fmo2*, *Fmo3*, *Fmo4* mRNA expression from normal lung tissue (n = 3), $EGFR^{T790M/L858R}$/CCSP ($E^{LT}$) mice tumors (n = 4), $KRAS^{G12V}$/$p53^{floxflox}$ (KP) mice tumors (n = 4) and EML4/ALK mice tumors (n = 4). Two-way ANOVA followed by Dunnet's multiple comparison. **b**) Representative images of FMO4 immunohistochemistry staining in human lung samples. On the left an example of non-tumoral lung tissue (normal lung), in the middle a case of FMO4-negative lung tumors, on the right an example of FMO4-positive case with expression in tumor cells. Scale bar: 250µm. **c)** FMO4 H-score in normal lung tissue (n = 244) and in the different lung cancer subtypes: lung adenocarcinoma (LUAD; n = 93), squamous cell carcinoma (LUSQ; n = 67) and other lung cancer subtypes (n = 25) from the same cohort as in **b**. Results are presented as violin plots, median and upper/lower quartiles are depicted with dashed or dotted lines respectively; **** $p$ <0.0001 (one-way ANOVA with Tukey's post hoc test). **d)** Frequencies of *FMO4* gene copy number alteration (CNA) in lung cancer samples from five different available databases. **e)** *FMO1*, *FMO2*, *FMO3* and *FMO4* expression levels in patients with lung cancer and 2 copies (diploid; n = 199 *for FMO1* and *FMO4* and n = 200 for *FMO2* and *FMO3*) or ≥ 3 copies (amp; n = 26 for all genes) of each gene locus. Z-scores (log RNA Seq V2 RSEM) were obtained from two studies (TCGA and PanCancer Atlas and OncoSG); * $p < 0.05$ (two-tailed unpaired Student's *t*-test). **f)** Kaplan Meier curves showing overall survival of lung cancer patients from the same cohort used in Fig 1d and Extended Fig. 1b,1c according to FMO4 expression (FMO4-low n = 48 and FMO4 high n = 135). Statistical significance was calculated with the log-rank (Mantel-Cox) test. # $p$ = 0.097 (HR = 1.505, 95% CI = 0.928–2.440).

**a** **b** **c**

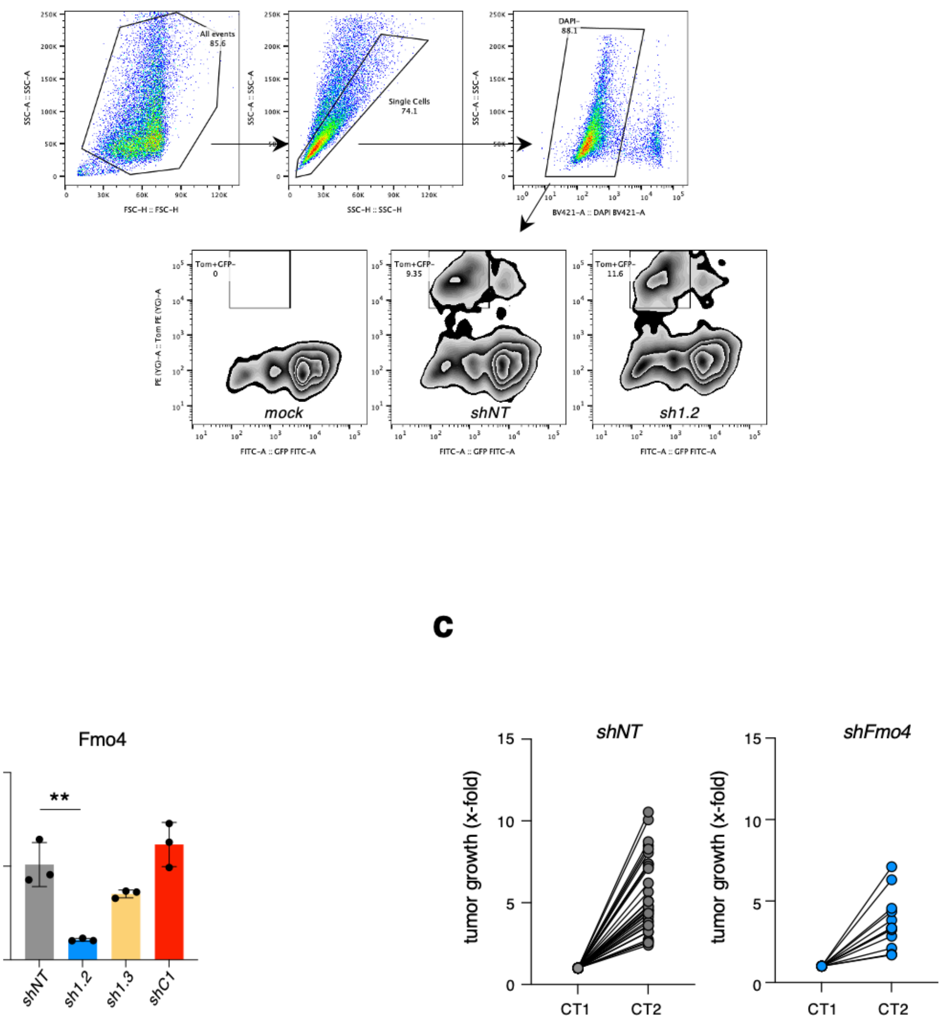


Extended Data Fig. 2, Bracquemond et al.

**Extended Fig. 2. FMO4 is needed for oncogenic KRAS-driven LUAD.**

**a)** Example of gating strategy for the sorting of M8-GFP-REV-Tomato murine cell line mock-infected (mock) or infected with lentiviral vector coding for Cre and a short hairpin RNA (shRNA) against FMO4 (*sh1.2*) or non-targeting (*shNT*) control. **b)** *Fmo4* mRNA expression from Tomato+-sorted M8 cells infected with lentiviral vector coding for Cre and a short hairpin RNA (shRNA) against FMO4 (*sh1.2*, *sh1.3*, *shC1*) or non-targeting (*shNT*) control. Results are presented as mean ± SD (n = 3); ** $p < 0.01$, (one-way ANOVA followed by Sidák's multiple comparison). **c)** Evolution of individual tumor volumes over 5 weeks, relative to the first measurement (CT1) in control shNT (n = 32 tumors) or shFMO4 (n = 11 tumors) in KP mice.

**a**

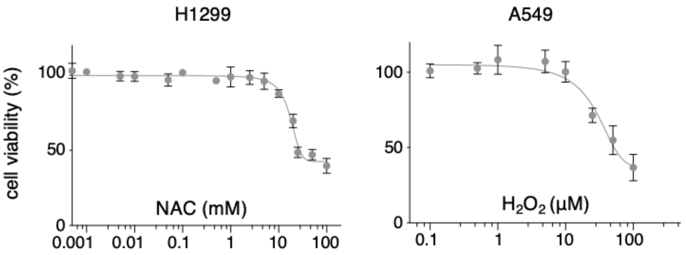


**b**

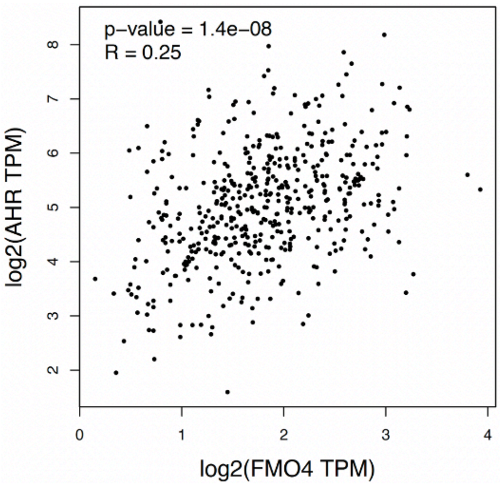


Extended Data Fig. 3, Bracquemond et al.

**Extended Fig. 3. Oxidative stress induces FMO4 expression through aryl hydrocarbon receptor and FMO4 loss of function increase reactive oxidative species.** **a**) On the left viability of H1299 cells incubated with the indicated NAC concentrations for 24 h. On the right viability of A549 cells incubated with the indicated $H_2O_2$ concentrations for 24 h. **b**) Correlation analysis between *AHR* and *FMO4* mRNA levels in human lung adenocarcinoma samples using the online tool http://gepia.cancer-pku.cn/detail.php?clicktag=correlation. The linear relationship was calculated using the Pearson correlation coefficient and was 0.25 (**** $p > 0.0001$).

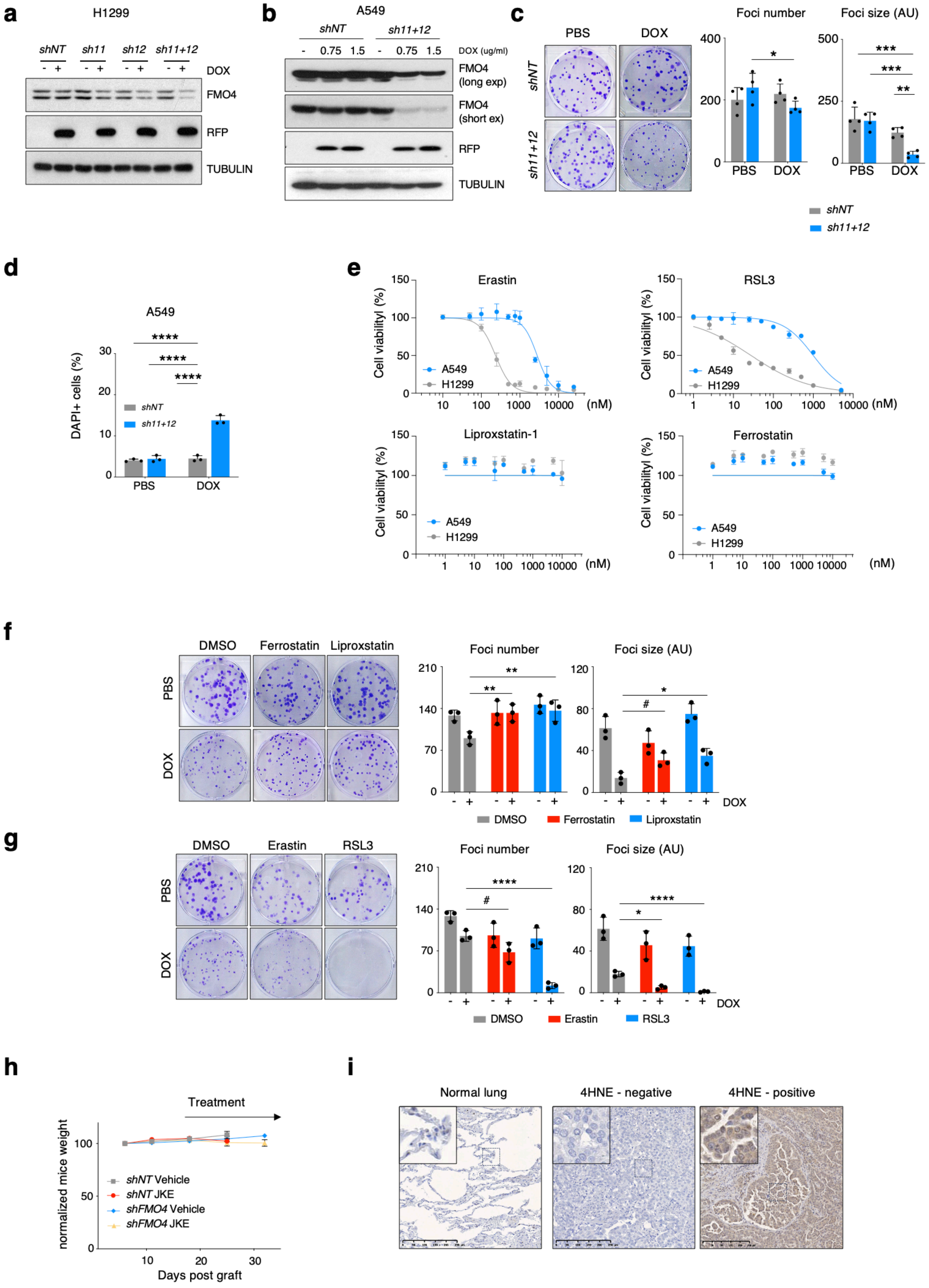


Extended Data Fig. 4, Bracquemond et al.

**Extended Fig. 4. FMO4 protect against ferroptosis in vitro and in vivo**

Immunoblotting of the indicated proteins in H1299 cells **a**) and A549 cells **b**) infected with inducible TRIPZ-shRNA lentiviruses targeting FMO4 (*sh11* or/and *sh12*) or non-targeting (*shNT*) control, and then incubated with 1μg/ml doxycycline (DOX) for 48 h. RFP was used as a reporter to monitor shRNA expression induction by doxycycline. **c**) Representative images of clonogenic assay in control (*shNT*) and *sh11+12*-infected A549 cells. Cells were plated at low density and incubated with PBS or doxycycline (DOX; 1.5 μg/ml) for 2 weeks. Quantification of foci number (left plot) and foci size (right plot) in each condition are presented as mean ± SD (n = 4); * $p < 0.05$, ** $p < 0.01$, ***p < 0.001, (two-way ANOVA with Fisher's LSD test). **d**) Quantification of the percentage of DAPI-positive staining in A549 cells by flow cytometry (n = 3); **** $p < 0.0001$ (two-way ANOVA with Fisher's LSD test). **e**) Viability of H1299 and A549 cell lines incubated with the indicated concentrations of ferroptosis activators (Erastin and RSL3) or inhibitors (Ferrostatin and Liproxstatin) for 72 h. **f)** Representative images of clonogenic assay in *shFMO4*-infected A549 cells incubated with PBS or doxycycline (DOX; 1.5 μg/ml), DMSO, Ferrostatin (5μM) or Liproxstatin (5μM), as indicated, for 2 weeks. Quantification of foci number (left plot) and foci size (right plot) in each condition are presented as mean ± SD (n = 3); # $p < 0.1$, * $p < 0.05$, ** $p < 0.01$, ***p < 0.001, (two-way ANOVA followed by Sidák's multiple comparison test). **g)** Representative images of clonogenic assay in *shFMO4*-infected A549 cells incubated with PBS or doxycycline (DOX; 1.5 μg/ml), DMSO, Erastin (100nM) or RSL3 (20nM), as indicated, for 2 weeks. Quantification of foci number (left plot) and foci size (right plot) in each condition are presented as mean ± SD (n = 3); # $p < 0.1$, * $p < 0.05$, ****p < 0.0001, (two-way ANOVA followed by Sidák's multiple comparison test). **h)** Change in mice body weight of mice grafted with H1299-*shNT* and -*shFMO4* cells. 18 days after tumor inoculation, mice were treated as in Fig. 4. Results are presented as mean ± SEM normalized to day 6 post-tumor engraftment. **h)** Representative images of 4-HNE immunohistochemistry staining in human lung samples. On the left an example of non-tumoral lung tissue (Normal lung), in the middle a case of 4-HNE-negative lung tumors, on the right an example of 4-HNE -positive case with expression in tumor cells. Scale bar: 250µm.

**a**

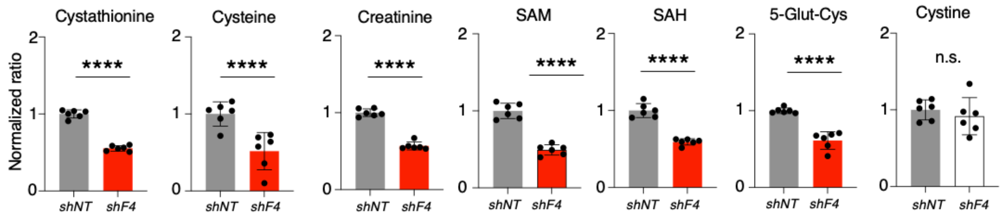


**b**

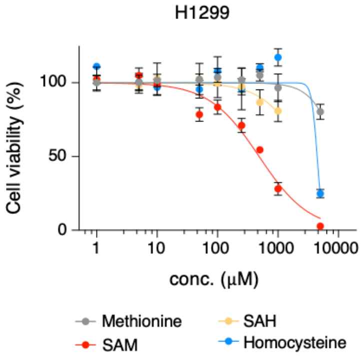


Extended Data Fig. 5, Bracquemond et al.

**Extended Fig. 5. FMO4 interacts with MAT2A and it is required for the proper function of the methionine cycle.**

**a)** Relative quantification of indicated metabolites in control (*shNT*) and *shFMO4*-transduced H1299 cells. Cells were incubated with doxycycline (DOX; 1 μg/ml) for 96h. Results are shown as mean ± SD (n = 6 each condition); * $p < 0.05$, (two-tailed unpaired Student's *t*-test). **b)** Viability of H1299 cells incubated with the indicated concentrations of Methionine, SAM, SAH or Homocysteine for 72 h.

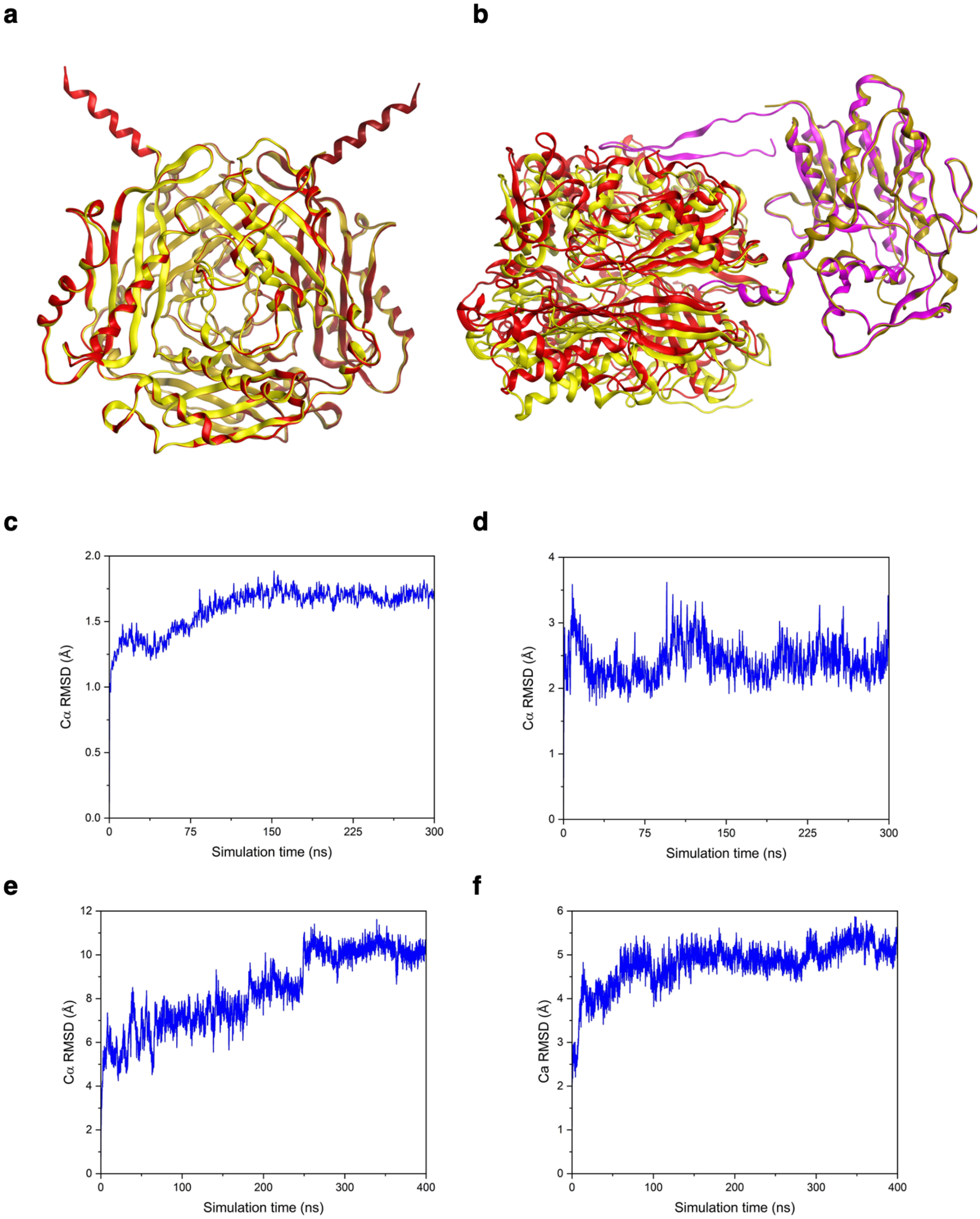


Extended Data Fig. 6, Bracquemond et al.

**Extended Fig. 6. FMO4 promotes the interaction between MAT2A and MAT2B.**

**a)** The AF3 model of $MAT2A_2$ (light and dark red color ribbons for each monomer) superposed on the Xray structure (yellow ribbon, pdb-id: 5A1I) resulted in a Cα RMSD of 0.348 Å. **b)** The AF3 model of $MAT2A_2$/MAT2B (red and magenta color ribbons for $MAT2A_2$ and MAT2B, respectively) superposed on the Xray structure (yellow ribbon, pdb-id: 4NDN) resulted in a Cα RMSD of 0.614 Å. The root mean square deviation (RMSD) plots of **c)** $MAT2A_2$, **d)** $MAT2A_2$/MAT2B, **e)** $FMO4_2$/$MAT2A_2$, and **f)** $FMO4_2$/$MAT2A_2$/MAT2B complexes during MD simulation courses.
